# Nucleoside diphosphate kinase A (NME1) catalyzes its own oligophosphorylation

**DOI:** 10.1101/2024.07.29.605581

**Authors:** Arif Celik, Felix Schöpf, Christian E. Stieger, Jeremy A. M. Morgan, Sarah Lampe, Max Ruwolt, Fan Liu, Christian P. R. Hackenberger, Daniel Roderer, Dorothea Fiedler

## Abstract

Protein phosphorylation is a central regulatory mechanism in eukaryotic cell signaling, and was recently expanded to include protein pyrophosphorylation and protein polyphosphorylation. Here, we report the discovery of yet another mode of phosphorylation – protein oligophosphorylation. Using site-specifically phosphorylated and pyrophosphorylated nucleoside diphosphate kinase A (NME1), the effects of these modifications on enzyme activity were investigated. Phosphorylation, and more so pyrophosphorylation, on threonine 94 notably reduced the nucleoside diphosphate kinase activity. Nevertheless, both phosphoprotein and pyrophosphoprotein were able to catalyze their own oligophosphorylation – up to the formation of a hexaphosphate chain – using ATP as a co-factor. This reaction was critically dependent on the catalytic histidine residue H118, and cryo-EM analysis of the differently modified proteins suggests an intramolecular phosphoryl transfer, likely *via* a phosphohistidine intermediate. Oligophosphorylation of NME1 in biochemical samples, as well as cell lysates, was further confirmed using mass spectrometry, and oligophophorylation promoted a new set of protein interactions. Our results highlight the complex nature of phosphoregulation, and the methods described here provide the opportunity to investigate the impact of this novel modification in the future.

## Introduction

Phosphorylation of proteins plays a central role in cellular signaling and contributes to the regulation of a plethora of processes.^1,2^ Following the characterization of serine, threonine, and tyrosine phosphorylation in mammalian cells, the investigation of protein phosphorylation has now been expanded to also include non-canonical amino acid side chains (histidine, arginine, cysteine, aspartate, glutamate, and lysine).^3–7^ Given the central involvement of phosphorylation in cellular signaling, a wide range of methods for the interrogation and characterization of protein phosphorylation have been developed over the past decades. For example, bottom-up phosphoproteomic analyses enable the annotation and comparison of thousands of phosphorylation sites on serine, threonine, and tyrosine side-chains in a single experiment.^8,9^ Highly selective pan-specific antibodies are available for the detection of phosphotyrosine (pY), and phosphoserine (pS) and phosphothreonine (pT) are readily detected within specific sequence.^10–12^ As a result, the protein data bank (PDB) contains an array of structures of proteins phosphorylated on serine, threonine, and tyrosine, which have contributed notably to our understanding of phosphoregulation.

By comparison, the tools for the investigation of non-canonical phosphorylation sites on histidine, arginine, cysteine, lysine, aspartate and glutamate are lagging behind. Useful antibodies for the detection of phosphohistidine (pHis) and phosphoarginine (pR) have been developed.^13–15^ But due to the higher lability of these phosphorylation sites, their characterization using mass-spectrometric and/or structural methods has been challenging.^3,7,16^

Another emerging mode of non-canonical phosphorylation is termed protein pyrophosphorylation, in which a pS or pT residue is further phosphorylated by inositol pyrophosphate messengers (PP-InsPs) to yield a diphosphorylated (or pyrophosphorylated) side chain.^17,18^ A recently developed mass spectrometry (MS) approach identified around 150 pyrophosphorylation sites, but structural characterization has been lacking completely so far.^19^ Moreover, the complexity of protein phosphorylation was further increased with the discovery of protein polyphosphorylation. This modification entails the attachment of inorganic polyphosphate (polyP) chains to PASK (polyacidic, serine-, and lysine-rich) domain-containing proteins.^20^ The exact nature of the polyP chain attachment is still under debate, and it is not clear if the polyP chain is covalently linked to lysine side chains *via* a phosphoramidate bond, or if the attachment is based on very strong non-covalent electrostatic interactions with the target sequences.^21–23^ Because structural and/or MS analyses of this intriguing modification have not been reported to date, is remains to be seen if one or both of these hypotheses can be validated.

A closer look at the pyrophosphorylation sites in human cells revealed that the large majority of sites reside in disordered, serine-rich polyacidic stretches.^19^ However, several pyrophosphorylation sites did not fit that pattern and localized to proline-directed consensus sequences within structurally resolved regions. Among these examples were a notable number of kinases, including glycogen synthase kinase-3 alpha/beta (GSK3 α/β), N-acetyl-D-glucosamine kinase (NAGK), and nucleoside diphosphate kinase A/B (NME1/2), and the pyrophosphorylation sites were close to the ATP-binding sites.^19^ In case of NAGK, it turned out that pyrophosphorylation was dependent on ATP (and not inositol pyrophosphates), and that the modification impacted the kinase activity as well as the protein interactions.^24^

NME1, best known as nucleoside diphosphate kinase A is a multifunctional enzyme that was found to be pyrophosphorylated on threonine 94 in human cells.^19^ A major function of NME1 is the synthesis of nucleoside triphosphates, other than ATP. NME1 catalyzes this process by transferring the γ-phosphoryl group from ATP to a nucleoside diphosphate (NDP) *via* an active-site phosphohistidine intermediate.^25^ Moreover, NME1 was recognized as a serine/threonine kinase and it phosphorylates the kinase suppressor of Ras 1 (KSR1).^26^ NME1 has also been proposed to be a mammalian histidine kinase.^27^ The transfer of the phosphoryl group from the phosphohistidine catalytic intermediate to histidine residues of protein substrates such as ATP-citrate lyase and succinate thiokinase could indeed be observed *in vitro* - the demonstration of NME1-mediated histidine phosphorylation in intact mammalian cells is an ongoing endeavor.^3,28–30^

For more than three decades, NME1 has been recognized for its role in suppressing metastasis in both melanoma and breast cancer, where NME1 expression levels directly correlate with its ability to suppress cell migration.^31,32^ However, phosphorylation at S120, S122 and S125 has been linked to tumor progression, highlighting the complex role of NME1 and its regulation *via* phosphorylation.^33,34^ Several additional phosphorylation sites on NME1 have been annotated in high-throughput studies (**Fig. 1a**), yet, the functional relevance of the most frequently detected phosphorylation site at T94, which is in close proximity to the ATP binding site, has not been investigated at all (**Fig. 1b**).^35^

**Fig. 1.**
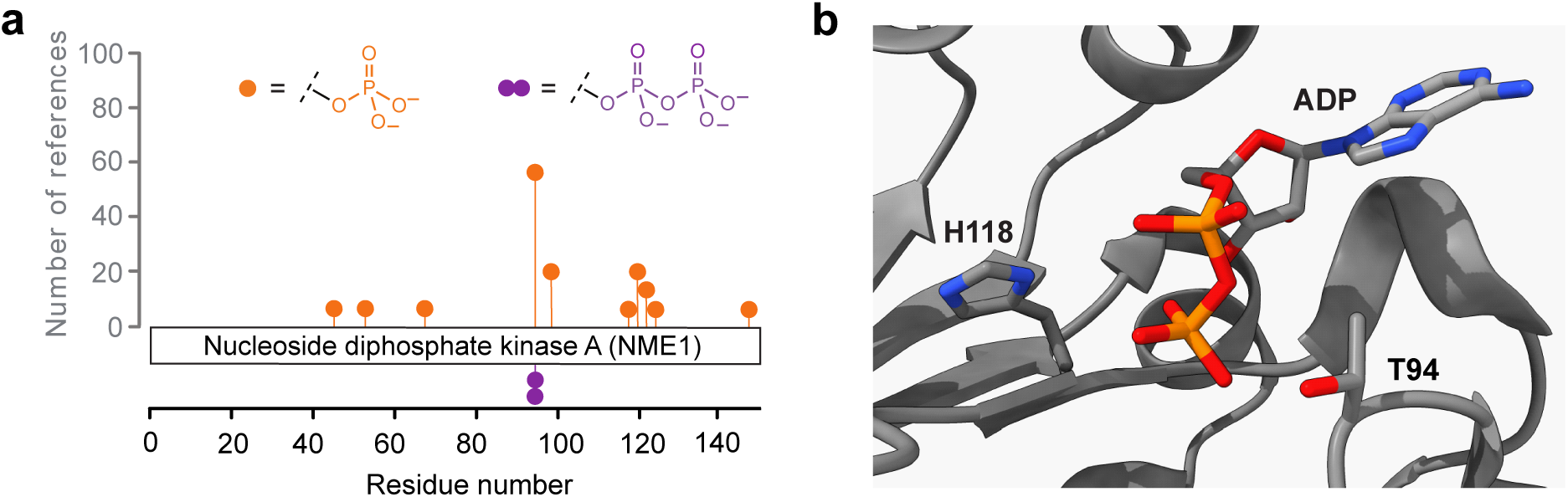
NME1 is phosphorylated and pyrophosphorylated near its substrate binding site. **a)** Phosphorylation sites on NME1 reported in PhosphoSitePlus^35^ (orange) and previously discovered pyrophosphorylation site at T94 (purple).^19^ **b)** Structure of NME1 co-crystallized with ADP, highlighting the location of T94, H118, and ADP (PDB code: 7ZLW). Carbon atoms are shown in grey, oxygen atoms in red, nitrogen atoms in blue, and phosphorous atoms in orange.

T94 is also the site at which NME1 is pyrophosphorylated, and we now report the synthesis and characterization of stoichiometrically phosphorylated and pyrophosphorylated NME1. Phosphorylation notably reduced NDP kinase activity and pyrophosphorylation led to complete inactivation of the enzyme. MS analysis and cryo-EM unveiled an ATP-dependent, autocatalytic formation of an oligophosphate chain on T94 *in vitro*, that only required the initial installment of a monophosphate group by cyclin dependent kinase 1 (CDK1). Oligophosphorylated NME1 was subsequently also detected in cell lysates, and this newly discovered post-translational modification appears to mediate a wide range of protein interactions. Our results highlight protein oligophosphorylation as yet another mode of phosphoregulation, and the methods described here will pave the way to investigate the occurrence and the function of this novel modification in the future.

## Results

### Generation of phosphorylated and pyrophosphorylated NME1

To investigate how phosphorylation and pyrophosphorylation at T94 may influence the properties of NME1, site-specifically and stoichiometrically modified protein is required. This was accomplished by employing a previously reported method that combines amber codon suppression and a chemoselective reaction using a photo-labile P-imidazolide reagent to obtain the protein in both phosphorylation modes.^24,36^ Initial attempts focused on the expression of pT94-NME1 in Escherichia coli (*E. coli*).^37^ Although the desired phosphoprotein pT94-NME1 was obtained in high purity, the yields were very low (0.05 mg/L, compared to 33 mg/L for wildtype-NME1) (**Supplementary Fig. S1 and S2**). Because a decent quantity of phosphoprotein is required for subsequent derivatization to the pyrophosphoprotein, we decided to incorporate pS as a surrogate for pT.^38^ This strategy proved highly effective, yielding high amounts of pS94-NME1 (0.75 mg/L) with the expected mass of 18168 Da (**Supplementary Fig. S3**). pS94-NME1 was then treated with biotin-polyethylenglycol-6-triazole-nitrophenylethyl-P-imdazolide (biotin-PEG_6_-Tz-NPE-P-imidazolide) overnight at 45 °C in a solvent mixture of DMA:H_2_O (9:1) (**Fig. 2a**). Following protein refolding, the formation of the derivatized protein, R-ppS94-NME1, was confirmed using quadrupole time-of-flight mass spectrometry (Q-TOF-MS). After exposure to 365 nm light, the pyrophosphoprotein was released, resulting in the isolation of ppS94-NME1. Compared to pS94-NME1, ppS94-NME1 displayed a mass increase of 80 Da, confirming the addition of a phosphoryl group (**Fig. 2b**).

**Fig. 2.**
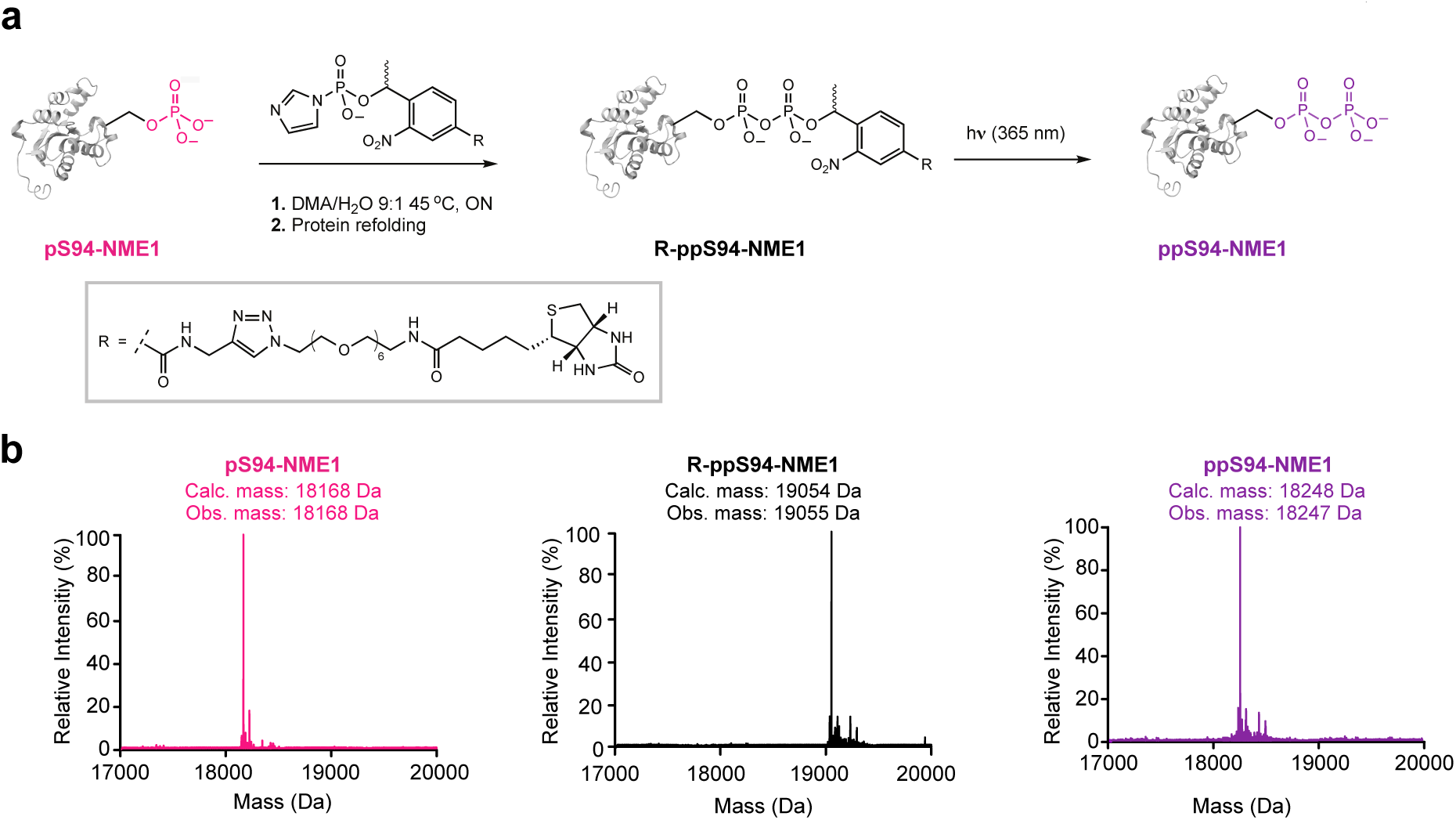
Expression and synthesis of pS94-NME1 and ppS94-NME1. **a)** Chemical phosphorylation of pS94-NME1 to provide pyrophosphoprotein ppS94-NME1. pS94-NME1 is derivatized by using the biotin-PEG_6_-Tz-NPE-P-imidazolide for the selective modification of phosphoserine moiety, to yield R-ppS94-NME1. Subsequent irradiation releases the pyrophosphoprotein ppS94-NME1. **b)** Deconvoluted Q-TOF-MS spectra of the intermediates and products of the reaction sequence in a) are shown.

To verify the effectiveness of the refolding step, the derivatization/refolding conditions were also applied to wt-NME1. The refolded protein displayed activity levels comparable to the reference protein (**Supplementary Fig. S4**), indicating successful restoration of the protein structure (**Supplementary Fig. S5**).

### Pyrophosphorylation reduces NME1 activity greatly

With the different (pyro)phosphoprotein variants of NME1 in hand, we next evaluated how these modifications impacted the NDP kinase activity of NME1, using a standard kinase assay with GDP or TDP as substrates and monitoring the consumption of ATP (**Fig. 3a**). With GDP as a substrate, wt-NME1 (at an enzyme concentration of 1 nM) exhibited high enzymatic activity (**Figure 3c**). By contrast, pT94-NME1 showed no detectable activity at the same enzyme concentration. Even when the enzyme concentration was elevated, the overall activity of pT94-NME1 was approximately 100-fold decreased compared to wt-NME1. The activity profile of pS94-NME1 was very similar to that of pT94-NME1, reduced approximately 100-fold compared to wt, suggesting that the pS94 mutation is a suitable substitute for pT94.

**Fig. 3.**
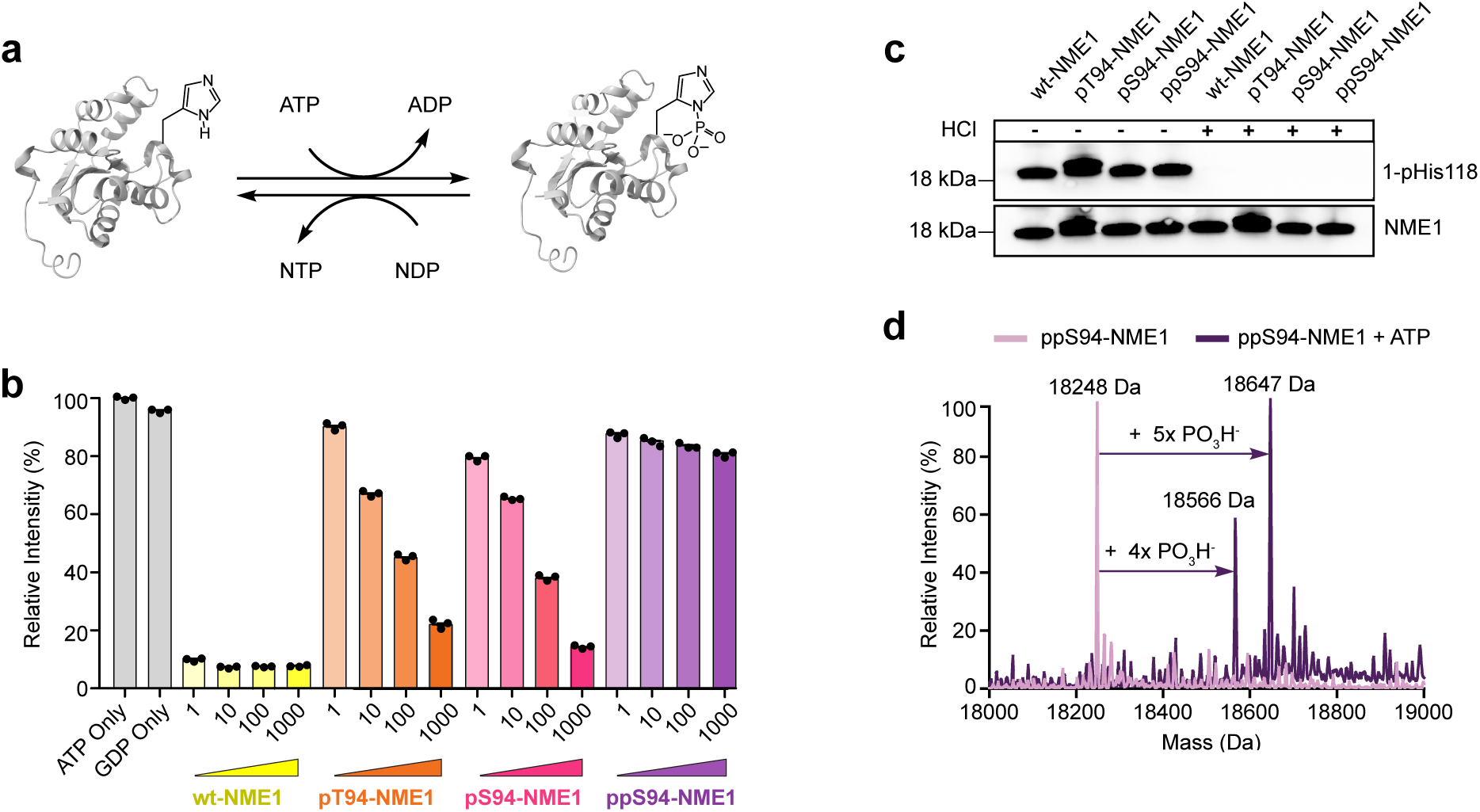
Phosphorylation and pyrophosphorylation of NME1 reduces the NDP kinase activity. **a)** NME1 catalyzes phosphoryl-transfer via an active site phosphohistidine intermediate. **b)** NDP kinase activity was measured utilizing GDP as a substrate, at 37 °C for 1 h in 50 mM Tris-HCl (pH 8.0), 150 mM NaCl, 10 mM MgCl_2_, 90 µM GDP, and 100 µM ATP. Concentrations of NME1 (1-1000) are in nM. Data is presented as mean ± SE of three technical replicates. **c)** Auto-phosphorylation reactions of NME1 were carried out at 37 °C for 1 h in 50 mM Tris-HCl, 10 mM MgCl_2_, 1 mM DTT, and 1 mM ATP. Hydrolysis was induced by incubating the auto-phosphorylated samples in 1M HCl for 1 h at 37 °C. Subsequently all samples were analyzed by western blot using anti-NME1 and anti-1-pHis antibodies.^14^ pT94-NME1 has a higher molecular weight due to the additional TEV cleavage site between the His_6_-tag and the NME1 sequence. **d)** ppS94-NME1 was incubated at 37 °C for 1 h in 50 mM Tris-HCl, 10 mM MgCl_2_, 1 mM DTT, and 1 mM ATP. Deconvoluted Q-TOF-MS spectra of ppS94-NME1 and ATP-treated ppS94-NME1 indicate additional phosphorylation.

The assay for ppS94-NME1 revealed that the pyrophosphoprotein exhibited almost no enzymatic activity at all, even at a protein concentration of 1 µM (**Fig. 3b**). A highly similar trend for the different (pyro)phosphoproteins was observed when TDP was used as a substrate: wt-NME1 demonstrated robust activity towards TDP, whereas the activity of pS94-NME1 was reduced approx. 100-fold; and ppS94-NME1 was essentially inactive (**Supplementary Fig. S6**).

Given that T94 is adjacent to the nucleotide binding site, phosphorylation or pyrophosphorylation at this site may make substrate access more difficult, due to electrostatic repulsion. The lowered NDP kinase activity of NME1 may therefore also correlate with reduced phosphorylation of the catalytic H118 residue, the critical residue for the phosphoryl transfer mechanism of NME1. However, for all (pyro)phospho-NME1 variants, the presence of the acid-labile 1-pHis118 intermediate was detected at similar levels using antibodies (**Fig. 3c**). The presence of an acid-labile phosphorylation was further confirmed by Q-TOF-MS for both wt-NME1 and pS94-NME1, following ATP treatment (**Supplementary Fig. S7**). Interestingly, when ppS94-NME1 was exposed to ATP, an addition of four or five phosphoryl groups (+360 and +400 Da) became apparent (**Fig. 3d**). This “hyperphosphorylation” was dependent on the catalytic H118 residue, as the kinase-inactive mutant ppS94-F118-NME1 did not undergo this modification (**Supplementary Fig. S8**).

Overall, the NDP kinase activity of NME1 is critically regulated by phosphorylation and pyrophosphorylation at T94/S94. Phosphorylation greatly reduced the kinase activity, and pyrophosphorylation virtually inactivates NME1. Surprisingly though, the pyrophosphoprotein underwent a hyperphosphorylation event when incubated with ATP, prompting us to further investigate this process.

### NME1 can undergo auto-oligophosphorylation

To understand in more detail where the additional phosphorylation events occurred when ppS94-NME1 is incubated with ATP, the reaction products were digested with trypsin and analyzed by tandem mass spectrometry (MS/MS) and label-free quantification (LFQ). Surprisingly, no additional phosphorylation sites could be identified, nor did any of the known phosphorylation sites show a significant increase following ATP treatment (**Fig. 4a**). The only notable change in the MS1 spectra was a large reduction of the signal for the pyrophosphopeptide upon exposure to ATP (**Fig. 4a**). Because the Q-TOF-MS clearly indicated the addition of several phosphoryl groups, we hypothesized that an extension of the pyrophosphate to an oligophosphate chain could explain the data.

**Fig. 4.**
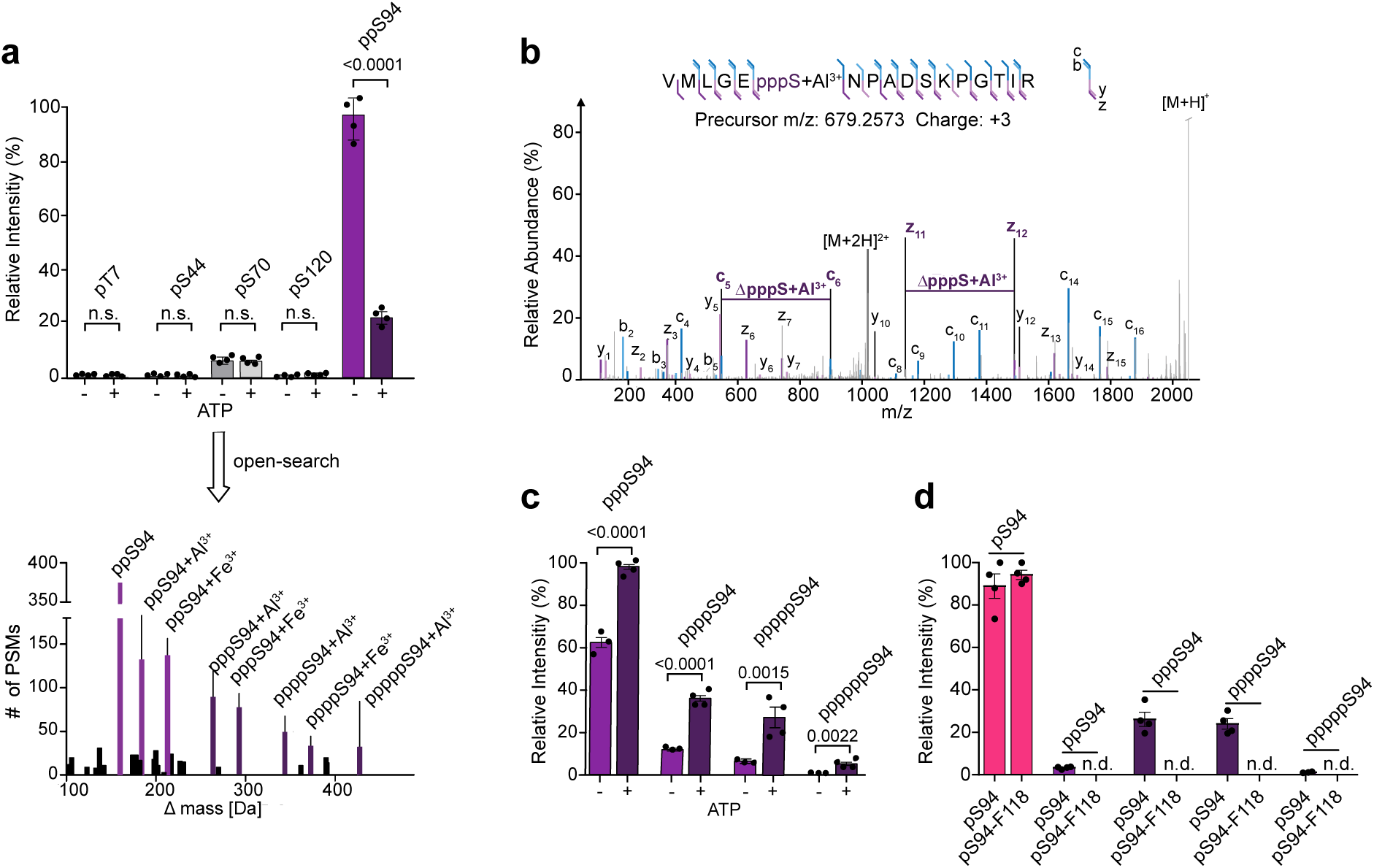
Characterization of oligophosphorylated NME1 using mass spectrometry. **a)** Relative quantification of ATP-treated ppS94-NME1 showed no increase of known phosphorylation sites but a decrease of pyrophosphorylation, while substantial number of oligophosphorylated PSMs revealed modification corresponding to oligophosphorylated NME1 peptides. n.s. = not significant and all samples were processed in technical replicates. **b)** Unambiguous localization of triphosphorylation on S94 by EThcD MS/MS. **c)** Relative quantification of different oligophosphorylation states before and after ATP treatment of ppS94-NME1. Assay was performed by incubating 1 µM ppS94-NME1 in 20 mM Tris-HCl (pH 8.0), 1 mM ATP, 5 mM MgCl_2_, 1 mM DTT. **d)** Relative quantification of different oligophosphorylation states in recombinantly expressed pS94-NME1 and pS94-F118-NME1. Data is presented as mean ± SE of four technical replicates.

To identify putative oligophosphorylation, the open-search feature integrated into MS Fragger was leveraged to facilitate an unbiased exploration of potential modifications.^39,40^ Alongside a substantial number of pyrophosphorylated peptide-spectrum-matches (PSMs; +160 Da, all localized to the sequence comprising S94), more than 100 spectra revealed a modification corresponding to the mass of pyrophosphorylation with the additional presence of Al^3+^ or Fe^3+^ ions, resulting in respective mass shifts of +184 Da or +213 Da (**Fig. 4a**). Remarkably, peptides that featured three to five phosphoryl groups on S94 were also identified. Further validation through manual inspection of the top-scoring spectrum-matches confirmed the precise localization of these modifications onto S94 (**Fig. 4b**).

Although this method enabled the unambiguous identification of oligophosphorylation on S94, many spectra exhibited poor fragment intensity. Instead, instances of electron transfer no dissociation (ETnoD) were observed, where the peptide ion loses a charge without undergoing subsequent fragmentation (**Supplementary Fig. S9**). Notably, this phenomenon was more pronounced in peptide ions containing an iron ion and could be attributed to the high electron affinity of Fe^3+^.^41^ Therefore, fragmentation techniques were screened and higher-energy C-trap dissociation (HCD) with low normalized collision energies (NCE 20-25) was identified as more reliable method to detect oligophosphorylated peptides **(Supplementary Fig. S10)**.^42,43^ As a result of optimization, a stepped-HCD method (NCE 20-23-26) was chosen and numerous tryptic peptides containing oligophosphate chains - up to hexaphosphate - were successfully identified, accompanied by excellent sequence coverage (**Supplementary Fig. S11**). Again, the peptides were exclusively observed as Fe^3+^-or Al^3+^-adducts, underscoring the strong chelating properties of oligophosphorylated peptides. Using relative quantification, a significant increase across all oligophosphorylation states was observed after ppS94-NME1 was treated with ATP (**Fig. 4c**).

Interestingly, oligophosphorylation was also detected in the ppS94-NME1 sample that had not been ATP-treated (**Fig. 4c**). Because it seemed unlikely that oligophosphorylation could occur during the chemical phosphorylation reaction, we reasoned that oligophosphorylation might have taken place prior to the derivatization - during the expression of the pS94-mutant in *E. coli*. To validate this hypothesis, pS94-NME1 was freshly expressed and analyzed by MS/MS, indeed revealing the autocatalytic formation of a pyrophosphate group at S94, as well as oligophosphate chains at this position. By contrast, neither wt-NME1, nor the double mutant pS94-F118-NME1 showed any higher phosphorylation states. These observations suggest an autocatalytic phosphoryl-transfer between H118 and pre-phosphorylated S94, to generate the pyro- and oligophosphorylation states. (**Fig. 4d**). Given that the innate modification site is a threonine residue, we also incubated pT94-NME1 with ATP and analyzed the sample via MS/MS using stepped-HCD, which clearly revealed pyro- and oligophosphorylation on threonine as well (**Supplementary Fig. S12**).

In summary, MS-based characterization of NME1 uncovered an autocatalytic pyrophosphorylation and oligophosphorylation mechanism *in vitro*, that critically depends on H118. Because pre-phosphorylation on T94 appears to be the only prerequisite for auto-oligophosphorylation, we wondered whether these higher modes of protein phosphorylation could also be observed within human cells.

### Oligophosphorylation of NME1 occurs in human cell lines

To potentially detect oligophosphorylated NME1 in complex samples, we adapted an enrichment workflow previously used for pyrophosphoproteomics (**Fig. 5a**).^19^ Briefly, HEK293T cells were lysed, digested and subsequently treated with λ-phosphatase to lower the amounts of monophosphorylated and multiply phosphorylated peptides. Given that the oligophosphorylated peptides have a high affinity for trivalent ions, a immobilized metal ion affinity chromatography (SIMAC) enrichment step was included to capture oligophosphorylated NME1 peptides. Following enrichment, peptides were eluted under basic conditions (0.5 % NH_4_OH) and examined using the optimized stepped-HCD method.

**Fig. 5.**
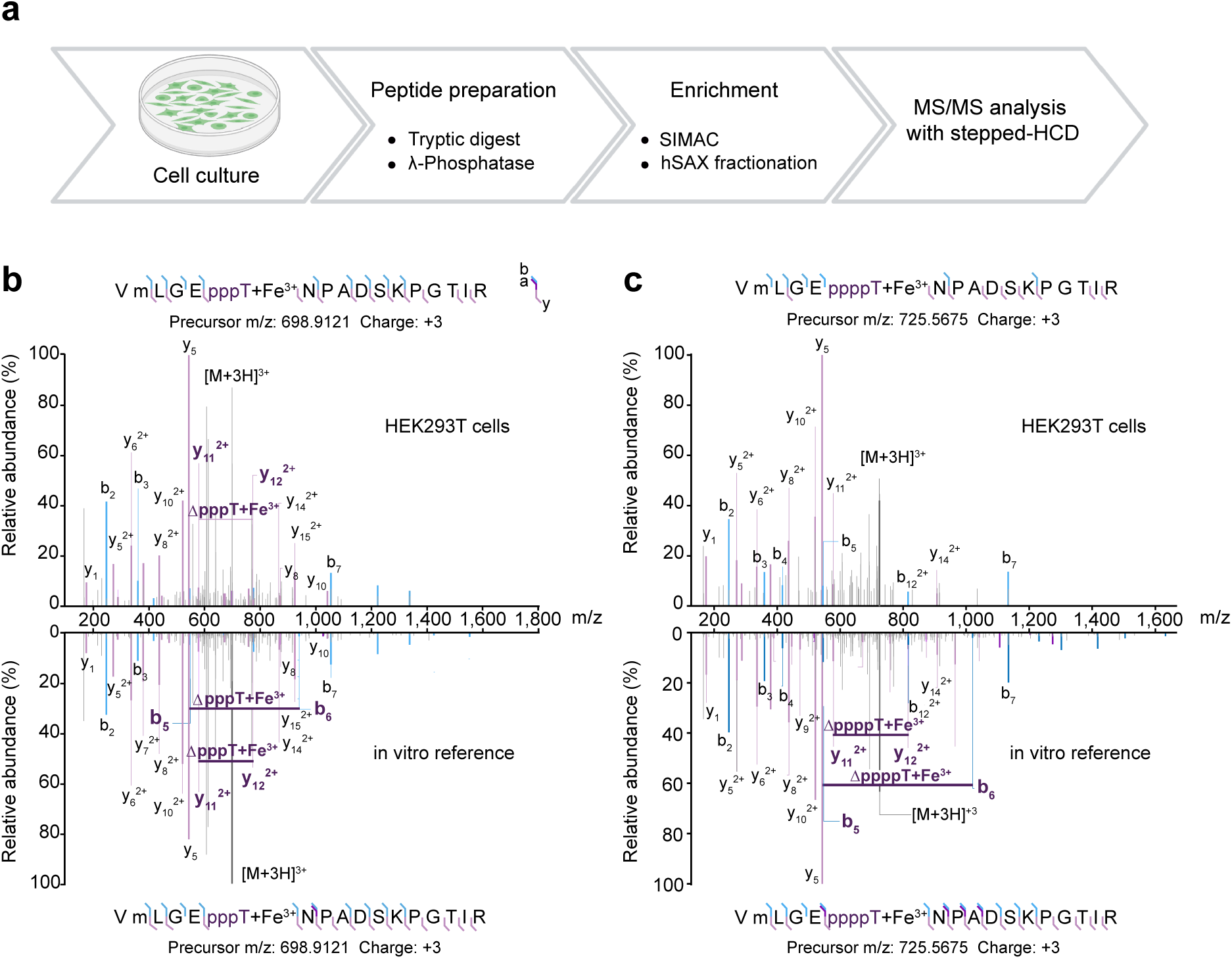
Enrichment and detection of oligophosphorylated NME1 peptides using MS/MS with stepped-HCD. **a)** Sample preparation workflow initially tailored to enrich tryptic pyrophosphopeptides. Comparison of stepped-HCD MS/MS spectra obtained from **b)** triphosphorylated and **c)** tetraphosphorylated NME1 peptide in HEK293T cells and recombinantly expressed pT94-NME1 shows excellent sequence coverage while preserving the modification.

Intriguingly, alongside the pyrophosphorylated NME1 tryptic peptide, the triphosphorylated and tetraphosphorylated NME1 peptides were readily identified. Ion couplets in the b/y series in the fragmentation spectra for the endogenous peptides (as well as the reference spectra) allowed the unambiguous localization of tri- and tetraphosphorylation of NME1 at T94, thereby confirming its occurrence in a human cell line (**Fig. 5b**). The measurements also substantiate the reliability of the stepped HCD-method for accurate identification of endogenously oligophosphorylated peptides, and will be a good starting point to develop bottom-up oligophosphoprotemic methods in the future.

### Phosphorylation by CDK1 reduces NDP kinase activity

Considering the pronounced effect of a single phosphorylation event on the NDP kinase activity, and the fact that phosphorylation of T94 is a prerequisite for pHis118 mediated oligophosphorylation, the question arose how this phosphorylation is installed. To date, no biochemical data on protein kinases targeting T94 on NME1 have been reported. However, a previous high-throughput proteomic investigation identified this residue as a potential site for phosphorylation by cyclin-dependent kinase 1 (CDK1).^44^ To investigate this connection, CDK1 was incubated with wt-NME1 in the presence of ATP and MgCl_2_. The addition of the phosphoryl group to NME1 was confirmed by Q-TOF-MS, and a significant increase of phosphorylation on T94 was measured by LFQ (**Fig. 6a,b**).^39^ At the biochemical level, the treatment of NME1 with CDK1 notably reduced NDP kinase activity, consistent with our previous observation using recombinantly expressed pS94-NME1 and pT94-NME1 (**Fig. 6c**).

**Fig. 6.**
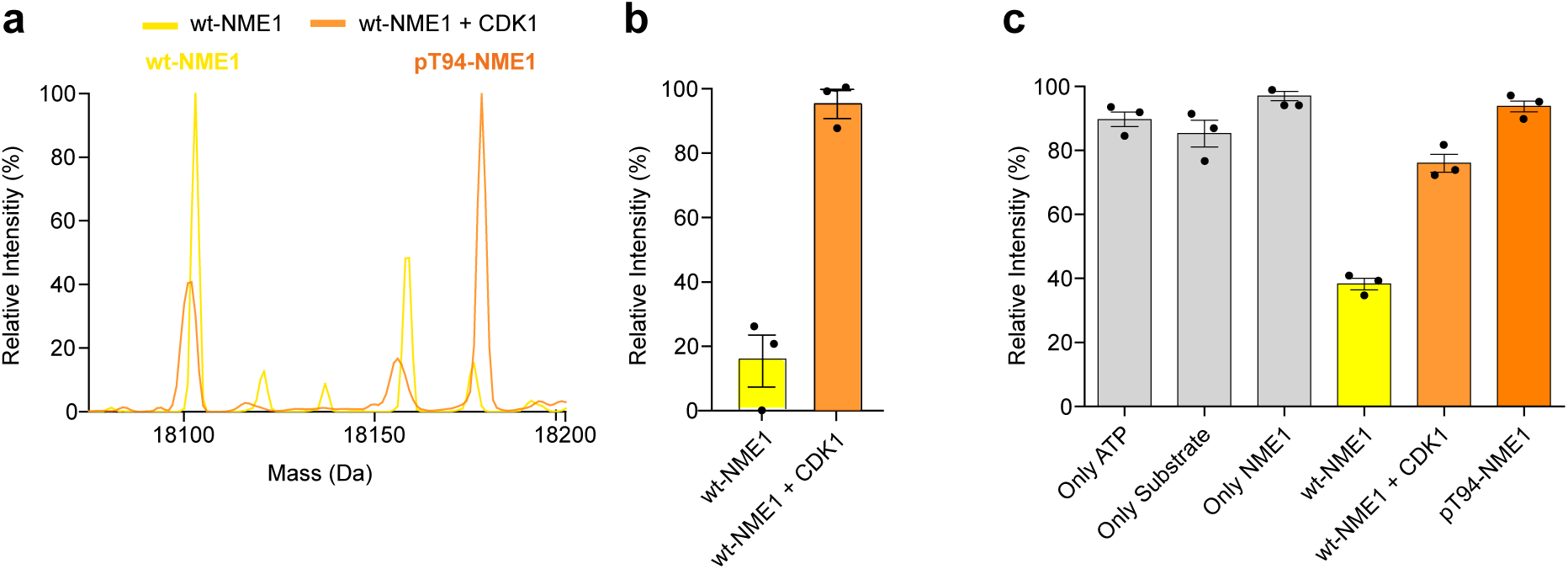
CDK1 catalyzes phosphorylation of NME1 on T94. **a)** Deconvoluted Q-TOF-MS spectra of wt-NME1 before (yellow) and after (orange) incubation with wt-NME1 shows the successful phosphorylation of wt-NME1 by CDK1. **b)** Label-free quantification (LFQ) of pT94 before and after CDK1 treatment showing the rel. intensity of pT94 **c)** NDPK activity of wt-NME1, before and after treatment with CDK1. The activity of pT94-NME1 is shown for comparison. Assays were performed by incubating 1 µM wt-NME1 and 50 nM CDK1 in 25 mM MOPS (pH 7.2), 10 mM MgCl_2_, 1 mM ATP, and 1 mM DTT at 37°C overnight. Data is presented as mean ± SE of three technical replicates.

Considering that the phosphorylation of NME1 by CDK1 can potentially instigate a whole sequence of phosphorylation events, and that the different phosphorylation modes regulate NDP kinase activity, we next evaluated the structural features of NME1 in different phosphorylated states

### Structure of pyrophosphorylated NME1 suggests intramolecular phosphoryl-transfer

We employed cryogenic electron microscopy (cryo-EM) and single particle analysis (SPA) to visualize potential structural alterations of NME1 following phosphorylation and pyrophosphorylation. For this, we vitrified purified pS94-NME1 and ppS94-NME1 and recorded cryo-EM datasets, which were refined to final resolutions of 2.8 Å for pS94-NME1, and 3.3 Å for ppS94-NME1 (**Supplementary Fig. S13 and S14**). Both density maps showed that the overall hexameric structure remained unaffected by the (pyro)phosphorylation, when compared to the wt-NME1 model (PDB: 7ZLW, **Fig. 7a**). Notably, additional cryo-EM density at S94 was observed in both cases, which corresponds to the incorporated monophosphate or pyrophosphate group. The resulting model for pS94-NME1 revealed that the phosphoryl group is directed towards the H118 side chain, with a distance of 6.6 Å between the monophosphate group and the 1-N of the H118 side chain (compared to 7.6 Å in wt-NME1). The model for ppS94-NME1 indicates a more pronounce effect, and the distance between the β-phosphoryl group and 1-N of H118 amounts to only 3.8 Å (**Fig. 7b**). Further efforts to visualize the link between both side chains by comparing the maps at the same threshold showed that the densities for the H118 side chain and the pyrophosphate group are forming a bridge in the ppS94-NME1 map (**Fig. 7c**). Thus, the short distance between H118 and ppS94 appears to be consistent with intramolecular auto-oligophosphorylation.

**Fig. 7.**
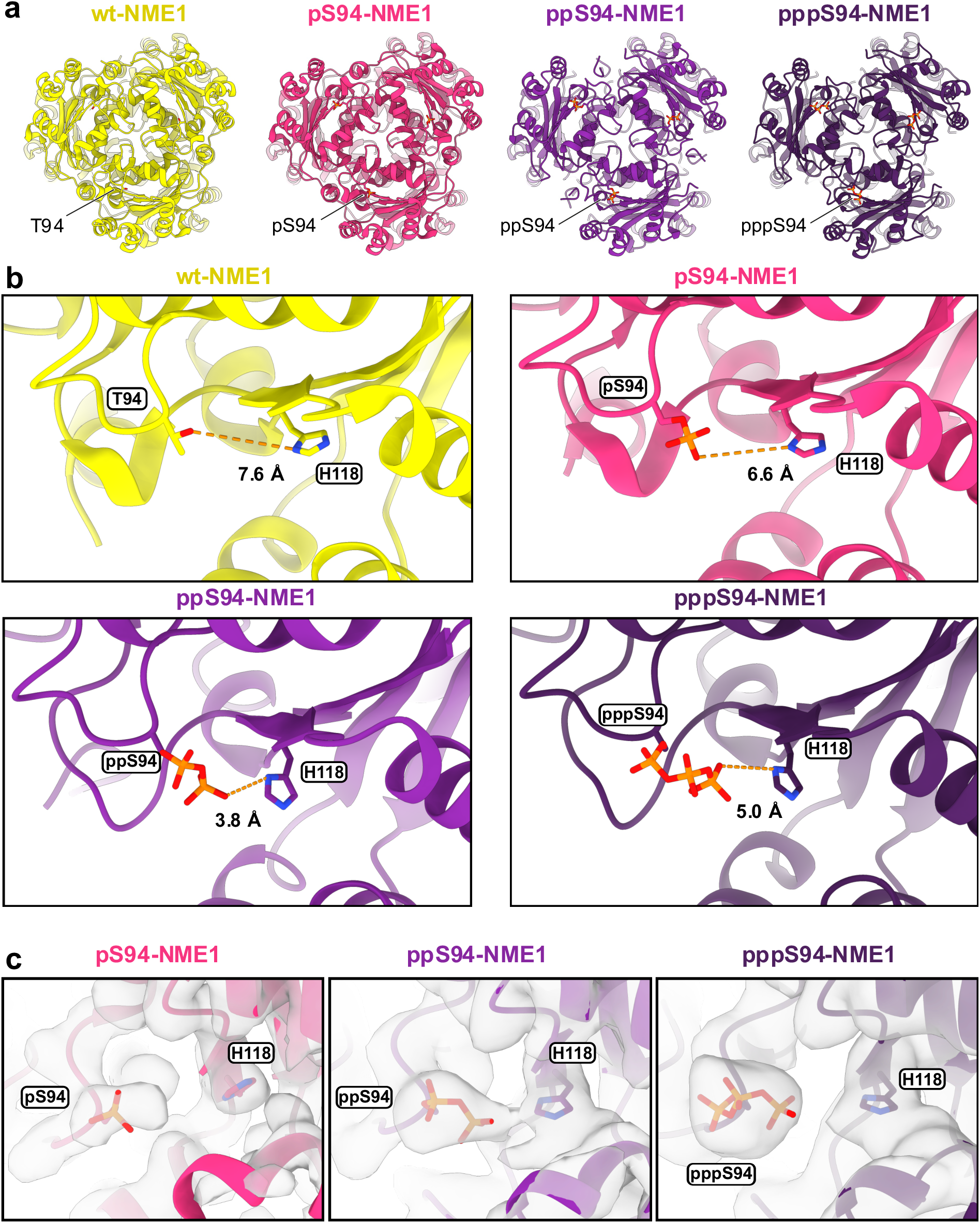
Cryo-EM structures of NME1 in different phosphorylation modes, as obtained at resolution of 2.8 Å for pS94-NME1, 3.3 Å for ppS94-NME1, and. 3.8 Å for oligo-pS94-NME1. **a)** Overview of the hexameric states of NME1 (wt - 7ZLW, pS94, ppS94, pppS94). Phosphoryl groups are highlighted. **b)** Distances between the terminal atoms of side chains of residue 94 (wt-T94, and phosphorylation modes on S94) and H118. **c)** Models of pS94, ppS94, and pppS94 in the corresponding cryo-EM maps plotted at similar thresholds, demonstrating the connecting density between ppS94 and H118.

Next, we sought to investigate the structural aspects of NME1 in oligophosphorylated states. Hence, purified ppS94-NME1 was treated with ATP to generate a heterogeneous mixture of oligophosphorylated NME1 for subsequent vitrification and cryo-EM data collection. After refining the data, a cryo-EM density map with an overall resolution of 3.8 Å was obtained (**Supplementary Fig. S15**).^45^ The hexameric structure still remained unaltered, even in these putatively oligophosphorylated states (**Fig. 7a**). While heterogeneity posed a challenge in visualizing individual oligophosphorylation states, we identified a larger density around ppS94. We were able to unambiguously verify that this density corresponds to the pppS94 species localized at S94. The distance between the γ-phosphoryl group and the 1-N of the H118 side chain now amounted to 5.0 Å (**Fig. 7b**). To visualize individual higher oligophosphorylation states in the heterogeneous data set, symmetry expansion and focused 3D classification of the phosphorylation site of one subunit were applied.^46^ This analysis revealed three subsets with different extents of cryo-EM density appearing around ppSer94, which is indicative of the presence of different chain-length oligophosphates (**Supplementary Fig. S16**). Class 1 contained 37 % of sorted particles and exhibited only little additional density, which likely corresponds to pppSer94 in a different orientation, whereas class 2 (32 % of particles) and class 3 (31 % of particles) had larger densities, possibly indicative of of higher phosphorylation states.

Furthermore, class 3 showed a connection between the additional density found for the oligophosphate chain and the density for the loop consisting of residues K65 - F61, which includes the positively charged R58 that could interact with the negatively charged oligophosphates.

Overall, this data presents the first high-resolution structural analysis of NME1 across higher phosphorylation states, demonstrating the reduced distance between pyrophosphorylated and phosphorylated S94 and the autocatalytic H118 side chain. The identification of additional cryo-EM density on the surface of the oligophosphorylated form by the subset class 2 and 3 may suggest a potential role in mediating protein-protein interactions and motivated us to identify such interactors.

### Many proteins preferentially interact with oligophosphorylated NME1

Given the known scaffolding role of inorganic polyP, as well as the observation that the protein polyphosphorylation of yeast nuclear localization sequence-binding protein (NSR1) negatively regulated its interaction with DNA topoisomerase 1 (TOP1), we next investigated whether oligphosphorylation of NME1 might alter its interactions with target proteins.^20,47,48^

For an initial reference data-set, HEK293T cell lysates were prepared and subsequently incubated with His_6_-tagged wt-NME1 or pS94-NME1 (in a buffer containing 150 mM NaCl to reduce non-specific ionic interactions). Following incubation, washing, and elution, the proteins were analyzed by bottom-up proteomics. We identified around 70 out of approximately 200 known interactors of NME1^49^; however, comparison of the LFQ values for pS94-NME1 versus wt-NME1 suggested that hardly any these protein interactions were affected by phosphorylation (**Supplementary Fig. S17**).

Next, the impact of oligophosphorylation on protein-protein interactions was explored by using oligo-pS94-NME1, which was generated by treating ppS94-NME1 with ATP. Again, while many known interactors could be identified, all of them remain largely unaffected by oligophosphorylation (**Supplementary Fig. S18**). The comparison of oligo-pS94-NME1 versus pS94-NME1, however, unveiled a unique set of protein-protein interactions that appeared to be dependent on oligophosphorylation. Specifically, around 80 proteins were identified that showed a prefential association with oligo-pS94-NME1, based on threshold set to log_2_ > 1.5 and -log_10_P > 1.5 (**Fig. 8a**).

**Fig. 8.**
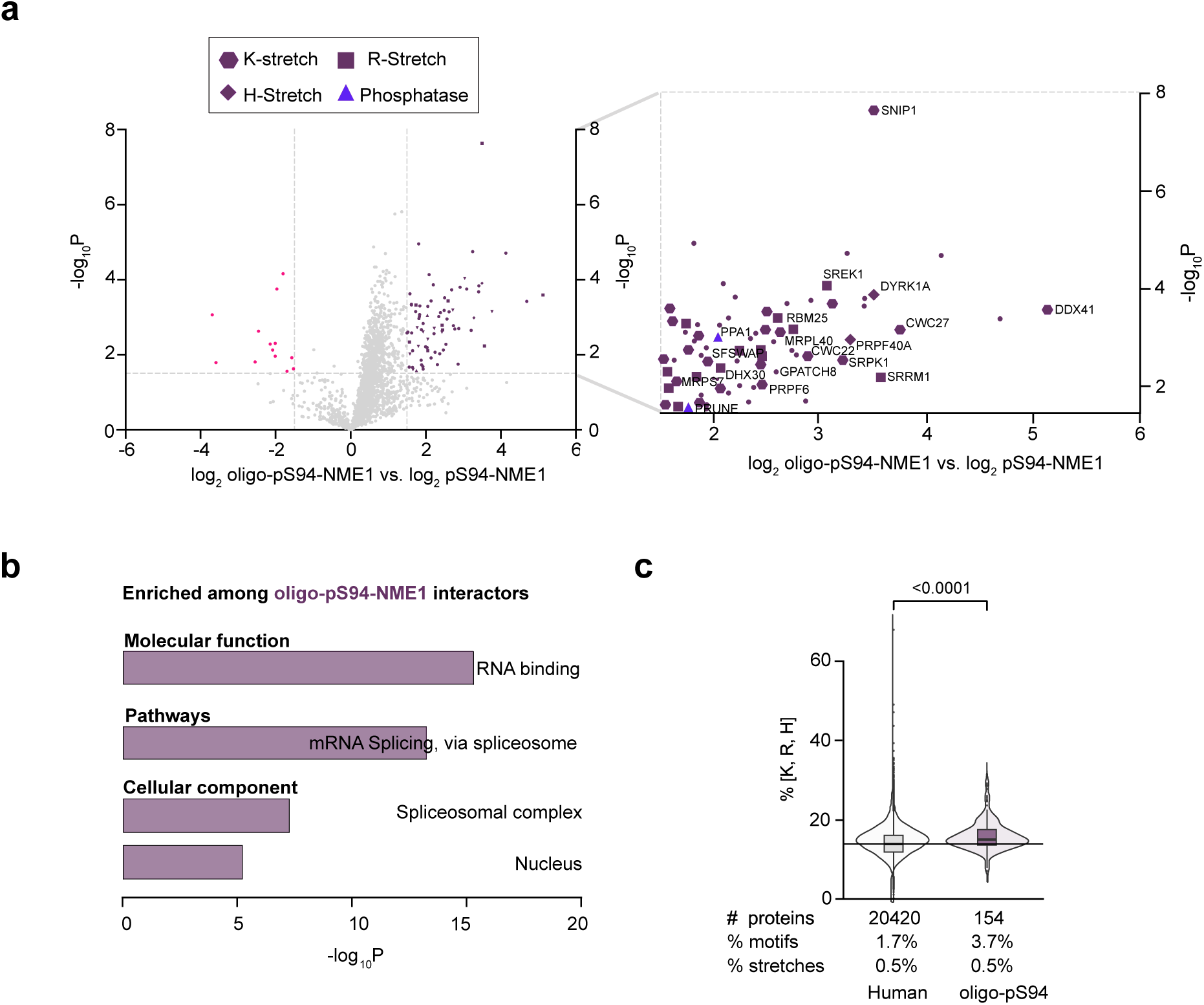
Interactome analysis of oligo-pS94-NME1 versus pS94-NME1. **a)** Volcano plot depicting LFQ values of oligo-pS94-NME1 versus pS94-NME1 after a t-test. The x-axis displays the difference of LFQ values on a log_2_ scale and the y-axis shows the -log_10_P value. The left side (pink) of the volcano plot illustrates the prefential enrichment with pS94-NME1; the right side (dark purple) illustrates prefential enrichment with oligo-pS94-NME1. Proteins with a log_2_ fold-change >1.5 and a log_10_P > 1.5 were considered to be significantly enriched. Named interactors have stretches of 4 or more H/K/R. **b)** Gene ontology analysis protein that prefentially interacted with oligo-pS94-NME1. **c)** Percentage of positively charged amino acids over total protein length for HEK293 proteome and oligo-pS94-NME1 interactors with t-test and a horizontal line highlighting the median percentage from the human proteome background. Below the sum of matched stretches or motifs divided by all proteins for each condition. Stretch: everything from 3 to 7 repeats of either K, R or H. Motif: triplett of either K, R or H allowing up to two interruptions.

To eliminate the possibility that the affinity enrichment strategy was strongly biased towards highly abundant proteins, the data was examined in relation to total protein abundance. Ranked proteins from a HEK293T whole-proteome study were plotted against their corresponding log10 iBAQ (intensity-based absolute quantification) values.^50^ The enriched proteins spanned an abundance range over seven orders of magnitude, including several proteins of low abundance, such as CWC22 (pre-mRNA-splicing factor CWC22 homolog) and SNIP1 (smad nuclear-interacting protein 1), but also high-abundance proteins were present (**Supplementary Fig. S19**).

To associate the proteins significantly enriched by oligo-pS94-NME1 with pathways or functions, we used Enrichr for gene ontology analysis **(Fig. 8b)**.^51,52^ This analysis suggests a role for oligo-pS94-NME1 in mRNA splicing *via* the spliceosome, as proteins from this pathway are heavily overrepresented among the interactors (CDC5L, PRPF40A, DDX41, CWC27, CWC22, SRRM1, SRPK1, SART1, PNN, PRPF6, TRA2B, SNIP1, RNPS1, PPIL3). Consistent with the over-representation of proteins involved in mRNA splicing is the association with the spliceosomal complex (cellular component) and RNA binding (molecular function).

Additionally, we observed an interaction between tyrosine phosphorylation-regulated kinase 1A (DYRK1A) and oligo-pS94-NME1. DYRK1A was recently shown to non-covalently bind to inorganic polyP *via* strong ionic interactions with polyhistidine stretches, which in turn negatively regulated the *in vitro* activity.^21^ As previous reports have showcased the ionic interactions with polyhistidine/polylysine stretches,^23^ we consequently screened the list of oligo-pS94-NME1 interactors for stretches containing consecutive H, K, or R residues and shorter interrupted H/K/R rich motifs, such as HxHH and HxxHH. Indeed, we found that the majority of the interactors harbored such positively charged motifs (**Fig. 8a**; **Table S1**).

Furthermore, we determined the percent composition of stretches and shorter interrupted motifs of all proteins prefentially enriched in the oligo-pS94-NME1 dataset and compared it to the HEK293 proteome. Interestingly, a significant enrichment of these motifs was present among the oligo-pS94-NME1 interacting proteins **(Fig. 8c)**. Finally, we identified two phosphatases, exopolyphosphatase PRUNE1 (PRUNE1) and inorganic pyrophosphatase 1 (PPA1), that exhibited a preferential interaction with oligo-pS94-NME1, suggesting that oligophosphorylation may be regulated by controlled removal of the oligophosphate chains by specific phosphatases.

## Discussion

NME1 continues to be a multifaceted enzyme with a diverse range of functions. We initially set out to investigate the influence of phosphorylation and pyrophosphorylation near the substrate binding site on the structure and function of NME1. To do so, genetic code expansion to produce stoichiometrically phosphorylated NME1 was combined with a chemical strategy to convert the phosphoprotein to the corresponding pyrophosphoprotein. When the NDP kinase activity of pS/pT94-NME1 and ppS94-NME1 was assessed, a strong, progressive reduction in activity was observed, compared to wt-NME1. Likely, the presence of the negatively charged (pyro)phophoryl group leads to significant electrostatic repulsion between the modification and the triphosphate moiety of ATP, slowing the rates of the formation of the pHis intermediate. Similary, subsequent binding of the nucleotide diphosphate will be slowed down, resulting in an overall decrease of catalytic activity. This observation is consistent with our previous findings where (pyro)phosphorylation close to the substrate binding site of N-acetyl-D-glucosamine kinase (NAGK) resulted in a strong reduction of enzymatic activity.^24^

As in the case of pyrophospho-NAGK, the formation of ppS94-NME1 was autocatalytic and ATP-dependent. Unexpectedly though, ppS94-NME1 proved competent to undergo subsequent ATP-dependent “hyperphosphorylation”, which turned out to be an auto-oligophosphorylation event. Employing MS/MS analysis with stepped-HCD, covalently attached oligophosphate - up to a hexaphosphate chain – could be detected and localized to S94. Formation of the pyro- and oligophosphate chains was dependent on the catalytic histidine residue H118, as the kinase inactive p/ppS94-F118-NME1 mutants lacked any signs of auto-oligophosphorylation.

Importantly, the stepped-HCD MS/MS analysis also uncovered the presence of oligophosphorylated NME1 in lysates from HEK293T cells, thus adding oligophosphorylation as a potential new mode of endogenous phosphoregulation. This observation then raises the question, whether oligophosphorylation is restricted to NME1, or if this modification occurs more broadly in eukaryotic cells. The initial MS/MS analysis already highlighted the complexities of detecting oligophosphorylated peptides – such as metal ion adduct formation – and warrants the development of a dedicated oligophospho-proteomics workflow. Such a development would greatly benefit from the availability of oligophosphorylated peptide standards, so that conditions for lysate preparation, peptide enrichment, and MS fragmentation methods can be systematically explored and optimized.

Our hypothesis that NME1 oligophosphorylation proceeds *via* an intramolecular mechanism is supported by the cryo-EM data we obtained for phosphorylated, pyrophosphorylated and oligophosphorylated NME1. Both the phosphorylated and pyrophosphorylated side chains are oriented towards H118, with a distance that would be compatible with an intramolecular transfer. As such, oligophosphorylation appears to result from a nucleophilic attack of the p/ppS94 residue on a 1-pHis118 intermediate. Interestingly, the structural data suggest, that the β-phosphoryl group of ppS94 is optimally placed (within 3.8 Å from H118) for a nucleophilic attack, and in closer proximity to the 1-pHis118 intermediated than the pS94 residue. This observation is corroborated by the biochemical data, which showed that ppS94-NME1 was much more competent to undergo oligophosphorylation, compared to pS94-NME1. In fact, when pS94-NME1 was treated with ATP, detection of the pyrophosphorylated intermediate was difficult, as it appeared to subsequently convert to the tri- and tetraphosphorylated species at a much higher rate. It will be interesting to investigate in the future, if certain stimuli, or added factors, can promote the “rate-limiting” addition of the β-phosphoryl group – thereby facilitating auto-oligophosphorylation.

Oligophosphorylated NME1 displayed a notable amount of cryo-EM density at the surface of the NME1 hexamer and it seems plausible that oligophosphorylation could act as a recruiting element to engage a different set of NME1 interaction partners. Indeed, many new interactors could be identified upon oligophosphorylation. The majority of these proteins harbored a stretch of polyhistidine, polylysine, or polyarginine and are predominantly associated with RNA splicing *via* the spliceosome. While the role of oligophosphorylation in these pathways remains to be determined, a simple explanation could be that the high negative charge of the oligophosphate modification resembles the negatively charged RNA backbone, thereby targeting a similar set of proteins.

The ability to catalyze the formation of oligophosphate chains is not an exclusive feature of NDP kinases. Many other enzymes across species are competent to catalyze phosphoanhydride bond formation.^53^ One notable set of enzymes encompasses the bacterial polyphosphate kinases PPK1 and PPK2, which are involved in the generation of long chains of inorganic polyphosphate.^47^ PPK2 is also known to have NDP kinase activity by catalyzing the transfer of phosphoryl groups from inorganic polyP to an NDP, while PPK1 catalyzes the synthesis of inorganic polyP from ATP using a pHis intermediate.^54–56^ Interestingly, these two functionalities seem to be united in NME1.

Given the remarkable reactivity of the pHis118-NME1 intermediate, the ability to generate oligophosphate chains may also be present in other human enzymes that harness a pHis intermediate. Interesting candidates include other proteins from the NME family (NME2-9), serine/threonine-protein phosphatase 5 (PGAM5), phosphoglycerate mutase 1 (PGAM1), 6-phosphofructo-2-kinase bisphosphatase (PFKFB3), as well as the kinase substrates of NME1, including succinyl CoA synthetase (SUCLG1), and ATP-citrate lyase (ACLY).^57–59^ For example, the crystal structure of PGAM5 illustrates the close proximity between the catalytic H105 and Y108 (5.6 Å).^57^ As Y108 is a known phosphorylation site, one could speculate that subsequent oligophosphorylation could take place (**Supplementary Fig. S20**).^60^

In principle, oligophosphorylation does not have to be an intramolecular process, but could also occur in trans. This may also be the case for NME1, which could – given that it is provided with the appropriate phosphoprotein substrate – act as an oligophosphate kinase. Therefore, much remains to be explored regarding the extent, the functional relevance, and the regulation of protein oligophosphorylation. All of the hypotheses and questions raised above will require the development of new analytical tools that can decipher, distinguish, and quantify all the different phosphorylation states of a given peptide or protein. The structural and mass spectrometric methods reported here provide the necessary starting points and should be built upon in the future.

## Supporting information

GO analysis

LFQ values for oligo-pS94-NME1 vs pS94-NME1 interactome analysis

LFQ values for oligo-pS94-NME1 vs wt-NME1 interactome analysis

LFQ values for pS94-NME1 vs wt-NME1 interactome analysis

Table for cryo-EM data collection, refinement and validation statistics

Table for raw data file designation

## Methods

### General information

All solvents and reagents were purchased from commercial sources and used without any further purification. All aqueous solutions were prepared using ultrapure laboratory grade water (deionized and filtered) obtained from an in-house water purification system. Antibiotics were prepared as stock solution at concentration of x1000 and stored at -20°C. Site-directed mutagenesis was performed on a Bio-Rad C1000 Touch Thermal Cycler machine. All bacterial growth media and cultures were handled using sterile conditions under an open flame. Protein concentrations were determined by the Pierce^TM^ Coomassie (Bradford) Protein-Assay-Kit (Thermo Fisher Scientific). Mass spectra were acquired on an Agilent 6220 TOF Accurate Mass coupled to Agilent 1200 LC (Agilent Technologies, USA). Intact proteins were analyzed using a Waters H-class instrument equipped with a quaternary solvent manager, a Waters sample manager-FTN, a Waters PDA detector, and a Waters column manager with an Acquity UPLC protein BEH C4 column (300 Å, 1.7 μm, 2.1 mm x 50 mm). LC-MS/MS analysis was performed using an UltiMate 3000 RSLC nano-LC system coupled on-line to an Orbitrap Fusion or Lumos mass spectrometer (Thermo Fischer Scientific). HEK293T cells were grown in 15-cm dishes to 70–80% confluency in Dulbecco’s Modified Eagle’s Medium (DMEM), supplemented with 10% FBS (Gibco), 1x Penicillin-Streptomycin (Gibco) and Glutamine (Gibco) (2 mM) in a 5% humidified CO_2_ incubator at 37°C. Cell counting was performed with the Bio-Rad TC-20 Cell counter. The following antibodies were used: Anti-N1-Phosphohistidine (1-pHis) Antibody, clone SC50-3 (Sigma-Aldrich), Anti-NME1 Antibody (Cell Signalling Technology), Anti-rabbit IgG, HRP-linked Antibody.

### Generation of expression plasmids

NME1 plasmid was obtained from GeneArt Gene Synthesis (Thermo Fischer Scientific) (pCDF-NME1 T94Ambercodon full-length), and the NME1 plasmid (pNIC28-Bsa-NME1) was a gift from the Kelly lab.

### Site-directed mutagenesis

Plasmid DNA harboring the corresponding NME1 ORF was extracted from an overnight culture of the pET21-NAGK vector in TB-Kan using a QIAGEN Miniprep kit. Single point mutations were installed by employing the Forward and reverse primer. 50 µl PCR reactions were performed on a Bio-Rad C1000 Touch Thermal Cycler following the NEB Phusion High-Fidelity DNA polymerase protocol using the following temperature program: 5 min 98 °C → 30 s 98 °C → 3 min 62 °C → Cycle to step 2 30× → 5 min 72 °C. The primers 5’- GCTGGGTGAGTAGAATCCTGCTG-3’ and 3’-ATCACGCGACCTGTCTTT-5’ were used to incorporate the amber codon at position 94.

The primers 5’- CAACATCATTTTTGGGTCGGACAG-3’ and 3’-CGGCCTACCTGAATACAG-5’ were used to replace H118 with F118.

### Expression and purification of His_6_-tagged wt-NME1

Chemically competent *E. coli* was transformed by heat-shock with plasmids containing wt-NME1 (pNIC28-Bsa-NME1). After recovery of 1mL of SOC medium for 1h at 37°C, the cells were cultured containing Kanamycin (50 μg/mL) and incubated overnight at 37°C. The overnight culture was diluted to an optical density of at 600 nm (OD_600_) of 0.05 in 1L of LB medium supplemented with Kanamycin (50 μg/mL) and cultured at 37°C, 400 rpm until OD_600_ of 0.6-0.8 was reached. IPTG was added to a final concentration of 1 mM, and protein expression was induced for 18h at 18°C, and Co-NTA affinity purification was performed using an FPLC system. After purification, the fractions containing the protein were collected, concentrated and rebuffered (50 mM Tris-HCl (pH 7.8), 150 mM NaCl, 1 mM DTT, and 10% Glycerol) using Amicon centrifugal filter units with a 10 kDa molecular weight cutoff (MWCO; Millipore)

### Expression and purification of His_6_-tagged pT94-NME1

Chemically competent *E. coli* BL21 (DE3) ΔserCΔycdX was co-transformed by heat-shock with plasmids containing both pCDF plasmid and pT94-NME1 (pET151a). After recovery of 1mL of SOC medium for 1h at 37°C, the cells were cultured containing Kanamycin (50 μg/mL) and Ampicillin (50 μg/mL) incubated overnight at 37°C. The overnight culture was diluted to an optical density of at 600 nm (OD_600_) of 0.05 in 4L of TB medium supplemented with Kanamycin (50 μg/mL) and Ampicillin (50 μg/mL) and cultured at 37°C, 400 rpm until OD_600_ of 0.6-0.8 was reached. IPTG was added to a final concentration of 1 mM and 2 mM (L)-O-phosphoserine (pH 7.0), and protein expression was induced for 18h at 18°C, and Co-NTA affinity purification was performed using an FPLC system. After purification, the fractions containing the protein were collected, concentrated and rebuffered (50 mM Tris-HCl (pH 7.8), 150 mM NaCl, 1 mM DTT, and 10% Glycerol) using Amicon centrifugal filter units with a 10 kDa molecular weight cutoff (MWCO; Millipore)

### Expression and purification of His_6_-tagged pS94-NME1 and pS94-F118-NME1

Electrocompetent *E. coli* BL21 (DE3) ΔserB was sequentially transformed by electroporation (Voltage: 1800 V, Capacitance: 25 μF, Resistance: 200 Ω, Gap length: 1.0 mm) with first pKW1-Sep (Camp^R^) and then with pNIC28-Bsa (Kan^R^) containing pS94-NME1 or pS94-F118-NME1. After recovery of 1mL of SOC medium for 1h at 37°C, the cells were cultured containing Kanamycin (50 μg/mL) and Chloramphenicol (25 μg/mL) incubated overnight at 37°C. The overnight culture was diluted to an optical density of at 600 nm (OD_600_) of 0.05 in 4L of TB medium supplemented with Kanamycin (50 μg/mL) and Chloramphenicol (25 μg/mL) and cultured at 37°C, 400 rpm until OD_600_ of 0.6-0.8 was reached. IPTG was added to a final concentration of 1 mM and 2 mM (L)-O-phosphoserine (pH 7.0), and protein expression was induced for 18h at 18°C, and Co-NTA affinity purification was performed using an FPLC system. After purification, the fractions containing the protein were collected, concentrated and rebuffered (50 mM Tris-HCl (pH 7.8), 150 mM NaCl, 1 mM DTT, and 10% Glycerol) using Amicon centrifugal filter units with a 10 kDa molecular weight cutoff (MWCO; Millipore).

### Nucleoside diphosphate kinase activity assay

Purified NME1 (1 nM–1000 nM) was added to a reaction mixture containing 100 µM ATP, 50 mM Tris-HCl (pH 8.0), 150 mM NaCl, 10 mM MgCl_2_, and 90 µM GDP or TDP. After 1 h at 37°C, *Promega* Kinase-Glo Plus^®^ reagent was added and the luminescence read out with a *Tecan* Infinite M Plex reader using 100 ms exposure time after 10 min of equilibration. Luminescence signals were read out with a TECAN Infinite M Plex plate reader.

### NME1 in vitro autophosphorylation

In vitro autophosphorylation of purified NME1 (50 ng/μL) was performed by incubating in 1 mM ATP in TMD buffer (20 mM Tris-HCl (pH 8.0), 5 mM MgCl_2_, and 1 mM DTT) at 37°C for 1h. Reaction were stopped by addition of 5x sample buffer (5X = 10% SDS, 250 mM Tris-HCl (pH 8.8), 0.02% bromophenol blue, 50% glycerol, 50 mM EDTA, 500 mM DTT; pH 8.8) and analyzed by modified SDS-PAGE conditions and Q-TOF-MS.

### NME1 in vitro phosphorylation by CDK1

1 µM wt-NME1 and 50 nM CDK1 was incubated in 25 mM MOPS (pH 7.2), 10 mM MgCl_2_, 1 mM ATP, and 1 mM DTT at 37°C overnight and subsequently analyzed by Q-TOF-MS, MS/MS and the NDPK activity.

### Generation of pyrophosphorylated NME1

For the chemical pyrophosphorylation of pS94-NME1, the reported protocol was used by only substituting the refolding buffer (50 mM Tris-HCl (pH 8.5), 35 mM KCl, 0.3 mM GSSG, 3 mM GSH, 10 mM EDTA, 0.2% CHAPS).^24^

### Q-TOF-MS

High-resolution ESI-MS spectra were recorded on two different instruments: 1) Agilent 6220 TOF Accurate Mass coupled to an Agilent 1200 LC (Agilent Technologies, USA) and were measured at 35 °C between 100–2000 m/z. The used column was an Accucore RP-MS (30 x 2.1 mm; 2.6 μm particle size) eluted with a flow of 0.8 mL/min and the following gradient (A = H_2_O + 0.1% TFA, B = MeCN + 0.1% TFA), gradient: 5% B 0–0.2 min, 5–99% B 0.2–1.1 min, 99% B 1.1–2.5 min. 2) Agilent Technologies 6230 Accurate Mass TOF LC/MS linked to Agilent Technologies HPLC 1260 Series; Column: Thermo Accucore RP-MS; Particle Size: 2.6 µM Dimension: 30 x 2.1 mm. The following gradient was used: A = H_2_O + 0.1 % formic acid, B = MeCN + 0.1 % formic acid, 5% B 0.0–0.2 min, 5-99% B 0.2–1.1 min, 99% B 1.1–3.6 min, 5% B 3.6–4.9 min. Flow rate: 0.8 mL/min; UV-detection: 220 nm, 254 nm, 300 nm.

### In solution tryptic digestion

Lyophilized samples were resolubilized in 100 µl Buffer (50 mM TEAB, 6 M Urea) and diluted to 2 M Urea. Samples were subsequently reduced and alkylated with 5 mM TCEP and 20 mM Iodoacetamide (IAA) at 37 °C for 1 h. Trypsin was added at an enzyme-to-protein ration of 1:100 (w/w) to digest overnight at 37 °C. Trypsin was then quenched with 1% FA (endconcentration) and centrifuged for 10 min at 20000 x g. The supernatant was collected and desalted using Sep-PAK C18 cartridges and lyophilized. Peptides were quantified by using BCA quantification.

### Liquid Chromatography and mass spectrometry

LC-MS/MS analysis was performed using an UltiMate 3000 RSLC nano-LC system coupled on-line to an Orbitrap Fusion mass spectrometer (Thermo Fisher Scientific) with instrument control software v3.4. For sample loading a PepMap C-18 trap-column (Thermo Fisher Scientific) of 0.075 mm ID x 50 mm length, 3 μm particle size, and 100 Å pore size was used. The loading mobile phase A contained 1% acetonitrile and 0.05% TFA acid in water, and mobile phase B 0.05% TFA acid in acetonitrile. Reversed-phase separation was performed using a 50 cm analytical column (in-house packed with Poroshell 120 EC-C18, 2.7 μm, Agilent Technologies) with mobile phase A containing 0.1% formic acid in water, and mobile phase B 0.1% formic acid in acetonitrile. The gradient started with 4% buffer B reaching 40% buffer B, with a total run time of 80- or 120-min including column wash and equilibration. MS1 scans were performed in the orbitrap using the following settings: resolution 120,000; AGC target 400,000; scan range 375 – 1500 m/z; max. injection time 50 ms. In 2 s cycles, precursors with a charge of 2 – 4 were sent to MS2 and excluded from fragmentation for 20 s. Depending on the purpose of the experiment, MS2 scans were acquired in the orbitrap with the following settings: (1) isolation window 1.6 m/z, resolution 15,000, HCD 23% or 30%, auto scan range, 22 ms max. injection time, 50,000 AGC target or (2) isolation window 1.6 m/z, resolution 60,000, EThcD with SA energy 30%, scan range 200 – 3000 m/z, 500 ms max. injection time, 100,000 AGC target.

### Analytical characterization of oligophosphorylated NME1

The obtained raw data was analyzed using FragPipe (v21) using the built-in open-search workflow^1^ and NME1_T94S plus contaminants as search space. The following MSFragger^2^ settings were applied: Precursor mass tolerance: -50 to +600 Da; Fragment mass tolerance: +/-20 ppm; Mass calibration & parameter optimization enabled; Isotope error: 0; Enzyme: Trypsin (cuts after K & R, no cut before P) with 2 missed cleavages, Peptide length: 7-50 AA; Peptide mass range: 500-5000 Da; Variable modifications: Oxidation (M, +15.9949 Da, up to 3x), Acetylation (N-term, +42.0106 Da); Carbamidomethylation of C was set as fixed modification (+57.02146 Da). Validation was performed using PeptideProphet (using default settings) and ProteinProphet. A protein-level FDR of 50% was applied. Additionally, PTM-Shepard^3^ was enabled (using the default settings). Identified peptides containing a modified S94 were filtered for at least five PSMs of the corresponding modification mass over all replicates. The best scoring PSM of each modification was manually validates and blotted using interactive spectrum annotator.

### Optimizing the fragmentation of oligophosphorylated peptides

Based on the obtained modification masses, an inclusion list was generated and samples were reanalyzed using a targeted method. LC-MS/MS analysis was performed using an UltiMate 3000 RSLC nano LC system coupled on-line to an Orbitrap Fusion mass spectrometer (Thermo Fisher Scientific). For sample loading a PepMap C18 trap-column (Thermo Fischer Scientific) of 0.075 mm ID x 50 mm length, 3 μm particle size and 100 Å pore size was used. The loading mobile phase A contained 1% acetonitrile and 0.05% TFA acid in water, and mobile phase B 0.05% TFA acid in acetonitrile. Reversed-phase separation was performed using a 50 cm analytical column (in-house packed with Poroshell 120 EC-C18, 2.7μm, Agilent Technologies) with mobile phase A contained 0.1% formic acid in water, and mobile phase B 0.1% formic acid in acetonitrile using a 93 minutes gradient (4-5%B 0-8 minutes; 5-25%B in 8-74 minutes; 25-28%B 74-80 minutes; 28-31%B 80-86 minutes; 31-36%B 86-92 minutes; 36-40%B 92-95 minutes; 40-50%B 95-96 minutes; 50-80%B 96-101 minutes; 80%B 101-104 minutes; 80-4%B 104-104.1 minutes). Data was acquired using survey scans in a range of 380 – 1400 m/z with a resolution of 120,000 and an AGC target value of 4e5. If a feature with m/z & z matching a peptide from the inclusion list was observed, 20 independent fragmentation events were triggered (see scheme). Fragment spectra were recorded in the orbitrap mass analyzer using 30,000 resolution and a maximum injection time of 100 ms with an AGC target of 50,000. Dynamic exclusion of 30 s was enabled after two full fragmentation cycles. Raw-files were analyzed as described above and obtained PSMs were manually inspected for fragmentation efficiency and unambiguous site-localization.

### HEK293T cell culture and lysate processing

HEK293T cells were grown in 15-cm dishes to 70–80% confluency in Dulbecco’s Modified Eagle’s Medium (DMEM), complemented with 10% FBS, Penicillin-Streptomycin (100 U/mL) and Glutamine (2 mM). Cells were washed twice with ice-cold DPBS (10 mL) and lysed by sonication (IKA Labortechnik, U200S control, 0.5 cycles, 50% intensity, 5 rounds) in 50 mM TBS buffer. supplemented with phosphatase and protease inhibitors (Roche PhosStop^TM^ and cOmplete^TM^ EDTA-free protease inhibitor cocktail). The cells were scraped off, transferred to protein low-binding microcentrifuge tubes, and incubated on ice for 10 min. The lysate was then centrifuged at 4°C for 10 min at 17,900 x g. The supernatants were combined and lysate protein concentration was determined using Pierce^TM^ Coomassie (Bradford) protein-assay-kit.

### Identification of oligophosphorylated peptides from HEK293 cells

For the enrichment of oligophosphorylated NME1 peptides the pyrophosphoproteomics sample preparation workflow was performed as described by Morgan et. al.^19^ Raw data was analyzed using FragPipe (v21) using the built-in offset-search workflow^1^ and NME1_T94S plus contaminants, NME1 plus contaminants, NME2 plus contaminants or the human proteome plus contaminants as search space. The following MSFragger settings were applied: Precursor mass tolerance: +/-10 ppm; Fragment mass tolerance: +/-20 ppm; Mass calibration & parameter optimization enabled; Isotope error: 0/1; Enzyme: Trypsin (cuts after K & R, no cut before P) with 2 missed cleavages, Peptide length: 7-50 AA; Peptide mass range: 500-5000 Da; Variable modifications: Oxidation (M, +15.9949 Da, up to 3x), Acetylation (N-term, +42.0106 Da); Carbamidomethylation of C was set as fixed modification (+57.02146 Da).

Additionally, the following masses were considered as offset-mass: 0.0 101.947 103.9256 105.9816 122.973 132.8784 136.9884 141.923 159.933 173.9882 177.9424 181.9128 184.0552 197.8802 199.8884 201.901 209.0186 212.845 216.9552 226.8986 228.8408 230.8544 263.8566 269.8656 292.8104 343.823 361.8338 372.7767 389.8042 390.788 423.7912 452.7431 527.7141 585.6209 79.9666 93.9798 94.0186 95.9618 98.0492. Validation was performed using PeptideProphet (using default settings) and ProteinProphet. A protein-level FDR of 1% was applied. Additionally, PTM-Shepard^3^ was enabled (using the default settings).

### Affinity capture experiments for proteomic analysis

All steps were conducted at 4 °C. A suspension of Nickel-beads (50 µL) was washed three times with 1 mL Milli-Q H_2_O and three times with 1 mL TBS Buffer (50 mM Tris-HCl pH= 7.5, 150 mM NaCl.) supplemented with 2 mM MgCl_2_ and 2 mM MnCl_2_. Subsequently, 25 µg of recombinant NME1 (wildtype, T94pS, ATP-treated T94ppS) in 100 µL TBS Buffer was added to the beads and incubated with constant rotation for 1 h at 4 °C. Beads were centrifuged at 2000 x *g* and the supernatant was discarded. The beads were washed three times with 1 mL TBS buffer and 1 mg of HEK293T cell lysate in TBS buffer was added to the beads and incubated for 3 h at 4 °C under rotation. Upon this time, the beads were centrifuged at 2000 x *g*, the supernatant was discarded and then they were washed three times with 1 mL TBS buffer. Lastly, proteins were incubated with elution buffer (50 mM Tris-HCl (pH 7.5), 150 mM NaCl, 500 mM Imidazole) for 1 h at 4 °C under constant rotation. After centrifugation at 2000 x *g*, the supernatant was collected and lyophilized.

### Identification, quantification, and statistics of interactome data

Raw data were analyzed and processed using MaxQuant software version 1.6.2.6a. Analysis was done with standard settings. Search parameters included two missed cleavage sites, fixed cysteine carbamidomethyl modification, and variable modifications including methionine oxidation and N-terminal protein acetylation. The peptide mass tolerance was 4.5 ppm for MS scans and 20 ppm for MS/MS scans. The match between-runs option was enabled. Database search was performed using Andromeda against the Human UniProt/Swiss-Prot database with common contaminants. The false discovery rate (FDR) was set to 1% at both the peptide and protein levels. Protein quantification was done based on razor and unique peptides. Label-free quantification was enabled. Bioinformatic analysis was carried out using Perseus software version 1.6.7.0. Proteins were filtered to exclude reverse database hits, potential contaminants, and proteins only identified by site. Proteins were further filtered by rows, requiring a valid value for at least two proteins out of four technical replicates. Data was imputed using Perseus default parameters, 0.3 width, and 1.8 downshift. Volcano plots were generated based on the log_2_ fold-change and the -log_10_(p-value) derived from a t-test (number of randomizations: 250).

### Physicochemical properties assessment

Proteomics data analysis and visualization were conducted using R (RStudio v2024.04.2). Proteins enriched with oligo-pS94-NME1 (log_2_-fold > 1.5) were compared to all 20,421 human protein sequences retrieved on 2024-07-10. Metrices derived from the primary amino acid sequence included the number of aromatic and charged residues, as well as the net charge per residue. The hydrophobicity of proteins was calculated using the peptides package, based and the Kyte & Doolittle scale.^61^ Secondary structure predictions were made using DECIPHER package.^62^ Modified residues were retrieved from UniProt. Stretches of positively charged residues were identified as three to seven consecutive repeats of the same amino acid (K, H, or R). Additionally, positively charged motifs were identified when a positively charged amino acid appeared three times, interrupted by up to two other amino acids, such as KxKK or KKxK.

### Cryo-EM sample preparation and data acquisition

For pS94-NME1 and ppS94-NME1, purified protein was vitrified on Quantifoil 1.2/1.3 Cu 300 mesh grids at 0.9 mg/ml using a Vitrobot Mark IV set to a blot force of 0, blotting time of 3.0 s. For oligo-pS94-NME1, ppS94-NME1 was incubated at 1 mM concentration with ATP for 3 h at 37°C before vitrification as above. Micrographs for all three data sets were acquired using a FEI Titan Krios G3i microscope (Thermo Fisher Scientific) operated at 300 kV equipped with a Bioquantum K3 direct electron detector and energy filter (Gatan) running in CDS counting mode at a slit width of 20 eV and at a nominal magnification of 105,000×, giving a calibrated physical pixel size of 0.83 Å/px on the specimen level. EPU 2.12 was utilized for automated data acquisition with AFIS enabled. For pS94-NME1, movies were recorded for 2.0 s accumulating a total electron dose of 47.7 e^−^/Å^2^ fractionated into 50 frames. Nominal defocus values were between -1.1 and -2.6 µm. For ppS94-NME1, movies were recorded for 2.0 s resulting in a total electron dose of 44.6 e^−^/Å^2^ fractionated into 50 frames with nominal defocus values between -1.2 and -2.4. For NME1-94Ser-Oligo-P, movies were recorded for 2.0 s accumulating a total electron dose of 44.1 e^−^/Å^2^ distributed over 50 frames. Here, nominal defocus values were between -1.4 and -2.4.

### Data processing of pS94-NME1, ppS94-NME1 and oligo-pS94-NME1

All data processing steps were carried out using CryoSPARC and are outlined in **Supplementary Fig. S12-14**.^63^ Obtained movies were aligned using Patch Motion correction and CTF was determined using patch CTF estimation. After sorting out bad images, micrographs were selected for further processing. Particles were selected and extracted for initial 2D classification and generation of autopicking templates. Subsequent template-based autopicking using and particle curation identified many particles. After three rounds of 2D classification (70 online-EM interations), an initial 3D hetero refinement (3 classes), and duplicate removal, particles were re-extracted without binning and subjected to non-uniform refinement using D3 symmetry and a molecular map of PDB 5UI4 (lowpass filtered to 12 Å) as an initial model.^64^ After iterative rounds of global and local CTF refinement and local motion correction, particles were sorted in two successive rounds of 3D hetero refinement (4 classes each), resulting in a final class of particles.^65^ These were then further processed using non-uniform refinement, followed by reference-based motion correction, yielding 532,096 (for pS94-NME1), 486,617 (for ppS94-NME1) and 201,312 (oligo-pS94-NME1) particles. A final non-uniform refinement resulted in a resolution of 2.8 Å (for pS94-NME1), 3.3 Å (for ppS94-NME1) and 3.79 Å (for oligo-pS94-NME1) according to the gold-standard Fourier Shell Correlation (FSC) criterion. DeepEMhancer^66^ was applied for map sharpening.

Detailed description for the data processing can be found in the supporting information.

### Atomic modeling of pS94-NME1

The atomic model of human NME1 (PDB 5UI4) was used as starting model and the attached imidazole fluorosulfate group and water molecules were removed. The model was rigid-body fitted into the sharpened density map using UCSF ChimeraX^67^, manually adjusted in Coot^68^, where Thr94 was mutated to phosphoserine, and ISOLDE^69^, and then refined using real-space refinement in Phenix^70^ (**Supplementary Fig. S12G, H**). Cryo-EM data processing and model refinement statistics are summarized in Table S1.

### Atomic modeling of ppS94-NME1 and pppS94-NME1

The atomic model of pS94-NME1 (see above) was used as starting point. The model was first rigid-body fitted into the obtained sharpened density maps for ppS94-NME1and oligo-pS94-NME1, respectively, using UCSF ChimeraX^67^ and then manually adjusted in Coot. Geometry retraints for ppS and pppS were generated using phenix.elbow^71^, and the resulting cif files were manually changed from a ligand to an amino acid. The changed amino acids were incorporated in the models, manually adjusted in Coot, and then refined using real-space refinement in Phenix^70^ (**Supplementary Fig. S13 and S14G, H**). Cryo-EM data processing and model refinement statistics are summarized in Table S1.

## Data availability

Supporting figures and tables are available in the Supporting Information.

The mass spectrometry proteomics data have been deposited to the ProteomeXchange Consortium via the PRIDE partner repository with the dataset identifier PXD054175.^72^

## Acknowledgments

A.C. gratefully acknowledges funding from the DFG (Deutsche Forschunsgemeinschaft) under project number 469186007. C.E.S was supported by a PhD fellowship of the Studienstiftung des Deutschen Volkes. We acknowledge access to electron microscopic equipment at the Core Facility for cryo-Electron Microscopy (CFcryoEM) of the Charité - Universitätsmedizin Berlin supported by DFG (INST 335/588-1 FUGG) for cryo-EM data collection, and we thank Dr. Thiemo Sprink for recording the data. The authors thank Jeffrey W. Kelly (The Scripps Research Institute) for supplying the pNIC28-Bsa-NME1 plasmid. The authors are grateful to Lena von Oertzen for their support in cell culture and Heike Stephanowitz for their support in digesting and desalting proteins for MS analysis. Finally, we thank all members of the Fiedler group for the insightful discussions and valuable input.

## Author contributions

D.F and A.C conceptualized the work, designed the experiments and wrote the paper. A.C carried out experiments including site-directed mutagenesis, protein expression, synthesis of pyrophosphorylated NME1, all biochemical validation as well as data analysis. C.E.S screened different fragmentation techniques and performed MS/MS analysis of oligophosphorylated peptides. F.S and D.R carried out the cryo-EM studies. S.L contributed sample preparation workflow for oligophosphorylated peptides and LFQ analysis. J.A.M.M contributed to MS data analysis. M.R carried out the proteomic data analysis and visualization by using R. F.L. provided guidance for proteomics experiments. C.P.R.H provided guidance for MS/MS analysis of oligophosphorylated peptides. C.E.S, F.S and D.R. wrote specific sections of the paper. All authors have read and approved the manuscript.

