## Supporting Information for "Nucleoside diphosphate kinase A (NME1) catalyzes its own oligophosphorylation"

[1]: Leibniz-Forschungsinstitut für Molekulare Pharmakologie (FMP), Robert-Rössle-Straße 10, 13125 Berlin, Germany

[2]: Institut für Chemie, Humboldt-Universität zu Berlin, Brook-Taylor-Str. 2, 12489 Berlin, Germany

[#] Authors contributed equally

This PDF file includes

Supplementary Figures S1 to S20

Extended Methods

Q-TOF Spectra

#### Supplementary Figures

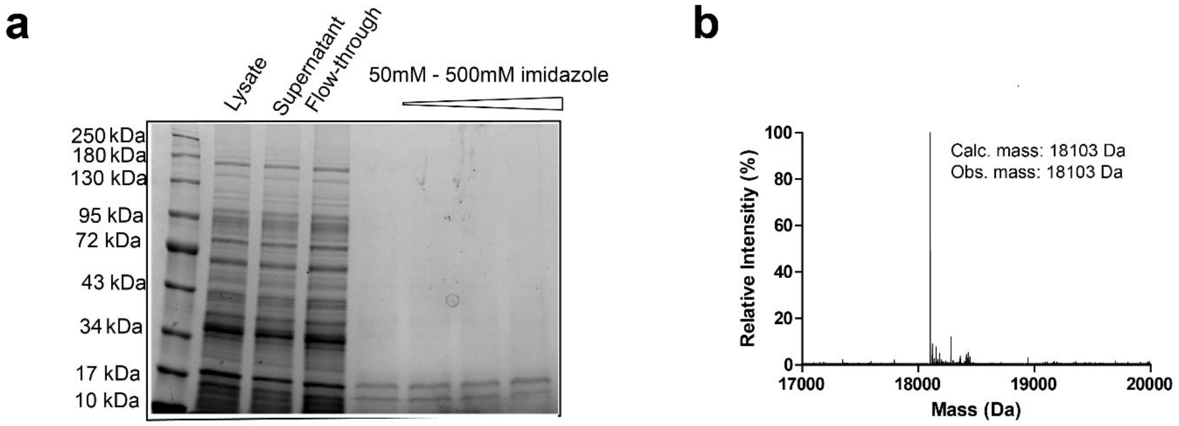


**Supplementary Figure 1.** Recombinant expression of wt-NME1. a) SDS-PAGE. Yield: 33 mg/L. b) Q-TOF-MS measurement.

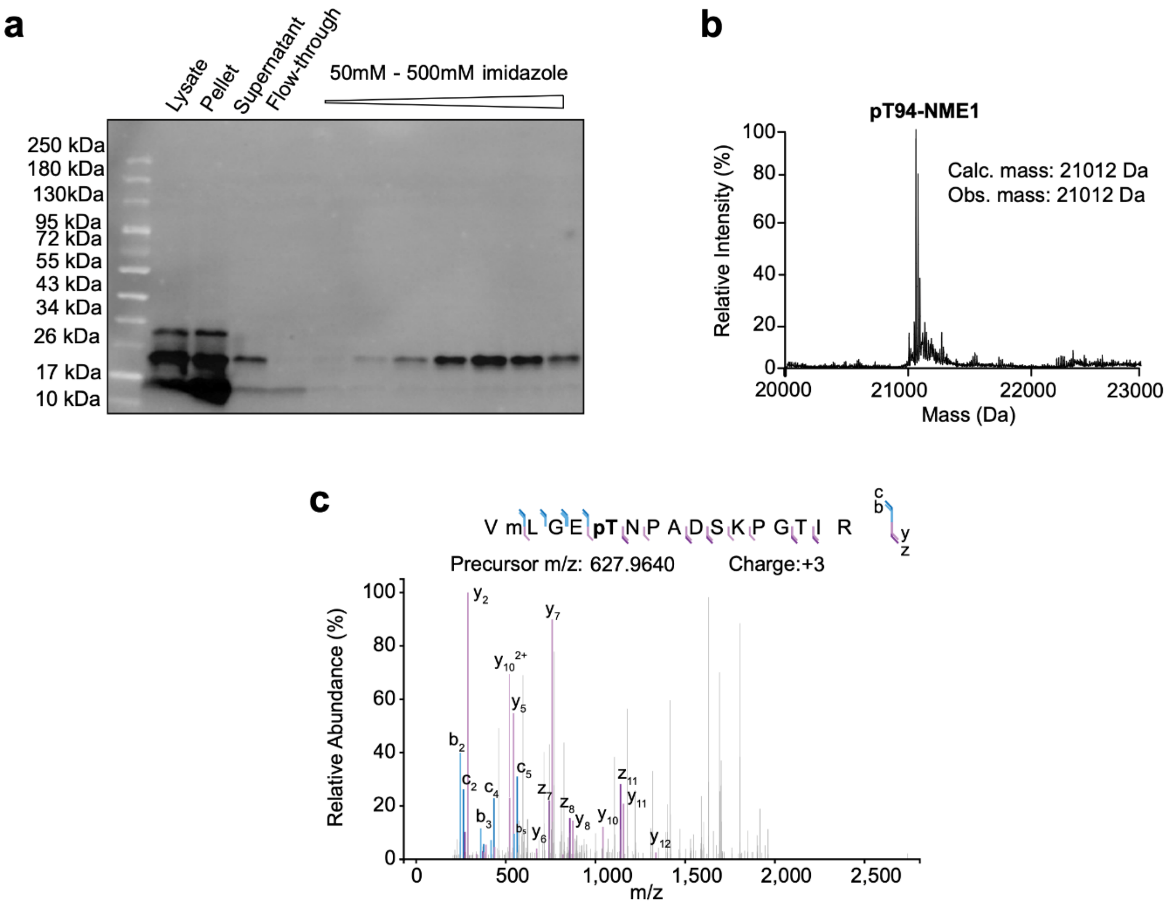


**Supplementary Figure 2.** Recombinant expression of pT94-NME1. a) SDS-PAGE. Yield: 0.05 mg/L. b) Q-TOF-MS measurement. c) MS/MS analysis localizing phosphorylation on T94.

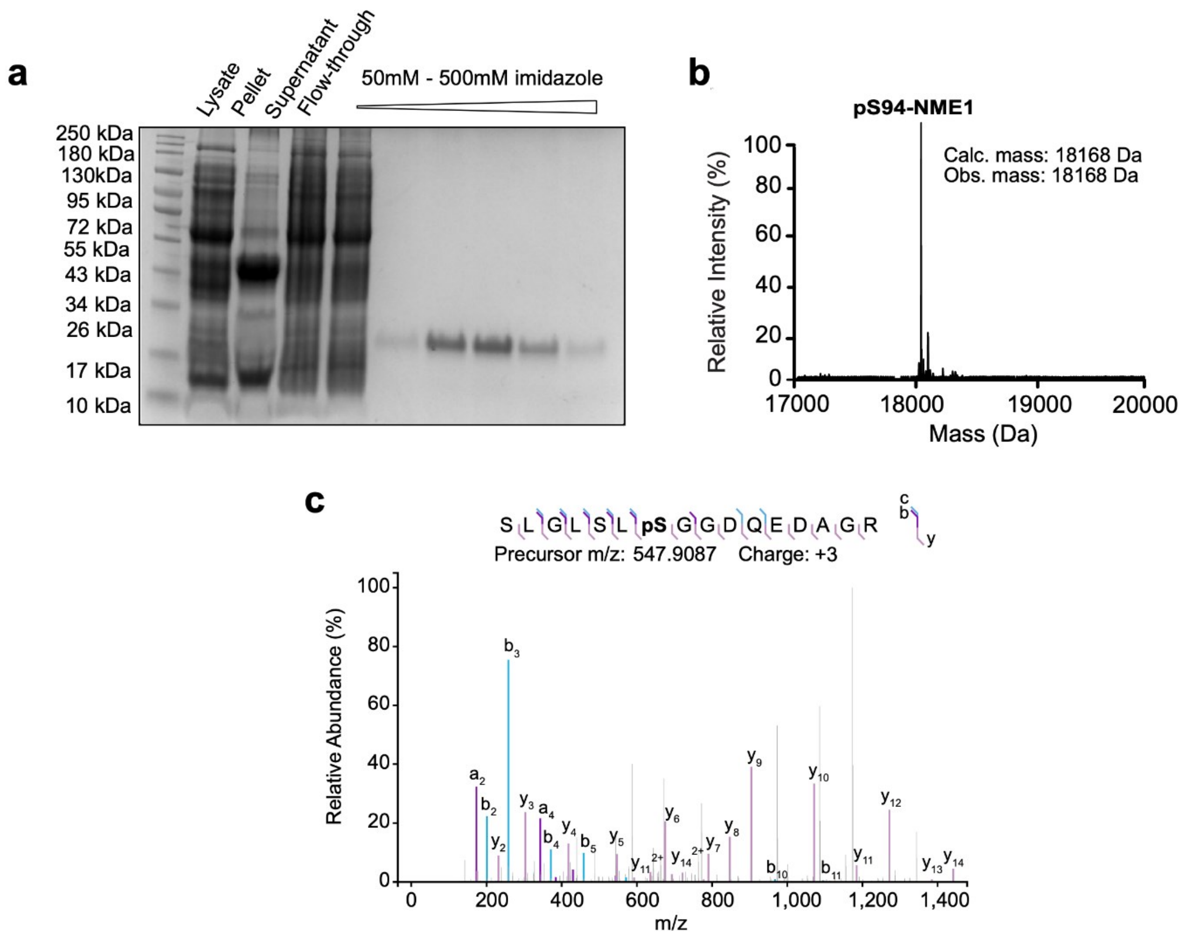


**Supplementary Figure 3.** Recombinant expression of pS94-NME1. a) SDS-PAGE. Yield: 0.75 mg/L b) Q-TOF-MS measurement. c) MS/MS analysis localizing phosphorylation on S94.

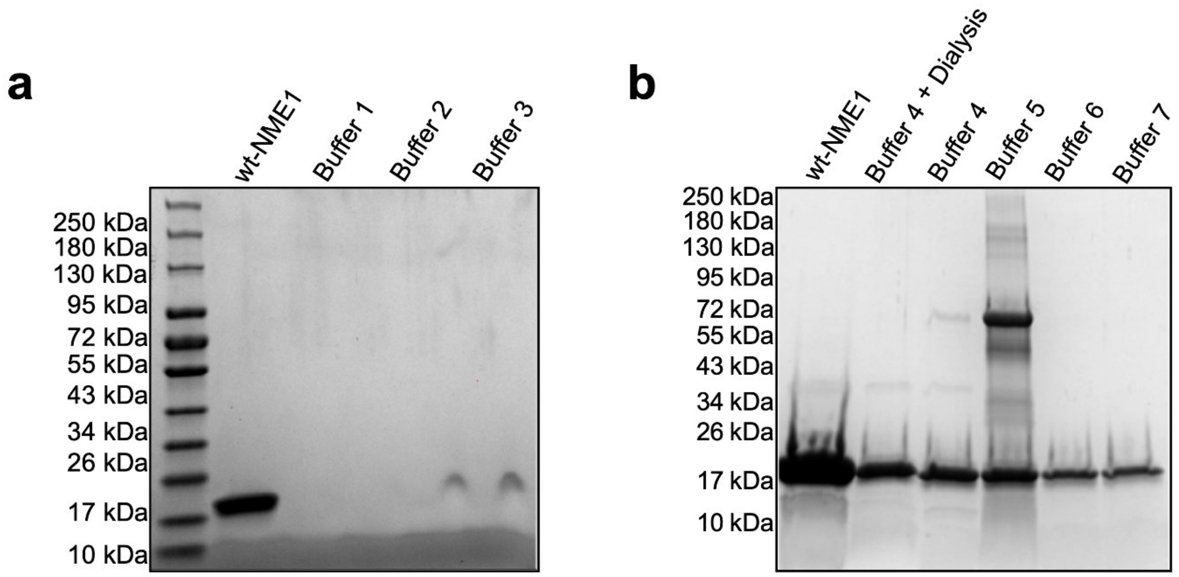


**Supplementary Figure 4.** Refolding of wt-NME1 analyzed by SDS-PAGE. a) SDS PAGE after applying the refolding protocol on wt-NME1. b) List of applied refolding buffers. **Buffer 2:** 50 mM Tris-HCl (pH 8.0), 35 mM KCl, 2 mM $MgCl_2$, 5 mM ADP, 100 mM L-Arg, 20 mM DTT. **Buffer 3:** 50 mM Tris-HCl (pH 8.0), 250 mM KCl, 2 mM $MgCl_2$, 5 mM ADP, 100 mM L-Arg, 20 mM DTT. **Buffer 4:** 50 mM Tris-HCl (pH 8.0), 500 mM NaCl, 50 mM $MgCl_2$, 50 mM L-Arg, 50 mM L-Glu, 5% glycerin. **Buffer 5**: 50 mM Tris-HCl (pH 8.5), 35 mM KCl, 0.3 mM GSSG, 3 mM GSH, 10 mM EDTA, 0.2% CHAPS. **Buffer 6**: 50 mM Tris-HCl (pH 8.5), 35 mM KCl, 20 mM DTT, 10 mM EDTA, 0.2% CHAPS. **Buffer 7**: 50 mM Tris-HCl (pH 8.5), 35 mM KCl, 0.3 mM GSSG, 3 mM GSH, 10 mM EDTA, 0.2% CHAPS, 400 mM Sucrose. **Buffer 8**: 50 mM Tris-HCl (pH 8.5), 35 mM KCl, 0.3 mM GSSG, 3 mM GSH, 10 mM EDTA, 0.2% CHAPS, 500 mM L-Arg.

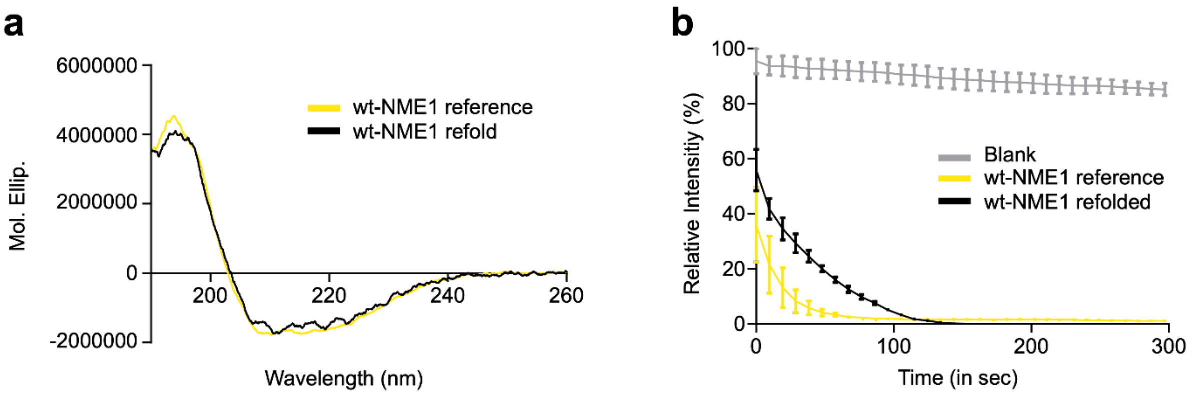


**Supplementary Figure 5.** Assessment of NAGK secondary structure and activity before after refolding. a) CD-Spectra of wt-NME1 before (in blue) and after refolding (in black). b) Nucleoside diphosphate kinase activity before and after refolding monitored by NAGH consumption. Assay condition: Purified NME1 (10 nM) was added to a reaction mixture containing 50 mM HEPES, 2 mM ATP, 2.2 mM ATP, 0.2 mM β-NADH, 1.1 mM PEP, 10 mM $MgCl_2$ 10 units lactic dehydrogenase and 7 units pyruvate kinase to a final volume of 100 µL. The reaction was vortexed immediately and a decrease in absorbance of NADH at 340 nm was then recorded for 5 min. UV signals were read out with a TECAN Infinite M Plex plate reader.

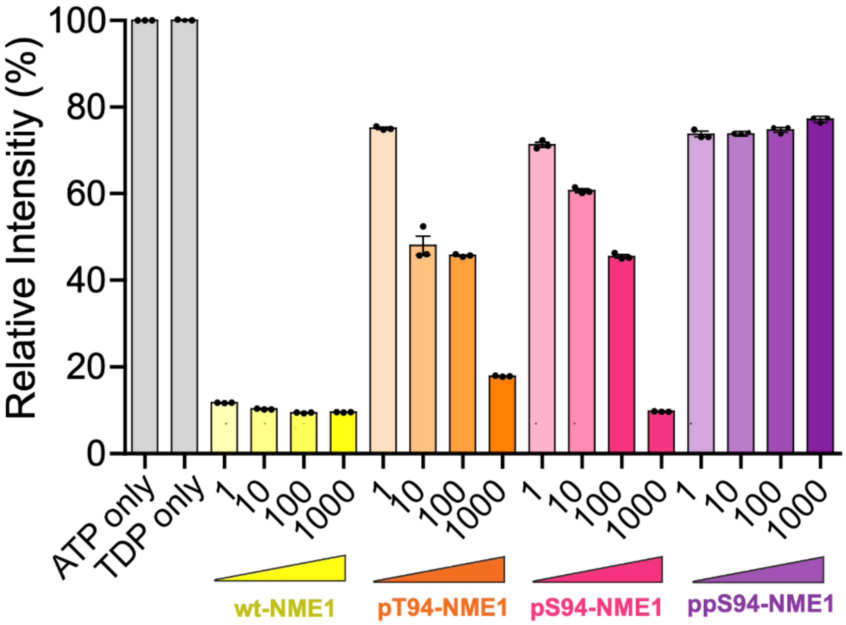


**Supplementary Figure 6.** Phosphorylation and pyrophosphorylation of NME1 reduces the nucleoside diphosphate (NDP) kinase activity. NDP kinase activity assay utilizing TDP as a substrate was measured at 37 °C for 1 h in 50 mM Tris-HCl (pH 8.0), 150 mM NaCl, 10 mM $MgCl_2$, 90 µM TDP, and 100 µM ATP. Concentrations of NME1 (1-1000) are shown in nM. c) Autophosphorylation of NME1 at 37 °C for 1 h in 50 mM Tris-HCl, 10 mM $MgCl_2$, 1 mM DTT, and 1 mM ATP with subsequent western blot analysis using anti-NME1 and anti-1-pHis antibodies. d) Deconvoluted Q-TOF-MS spectra of ATP-treated ppS94-NME1 indicating additional phosphorylation.

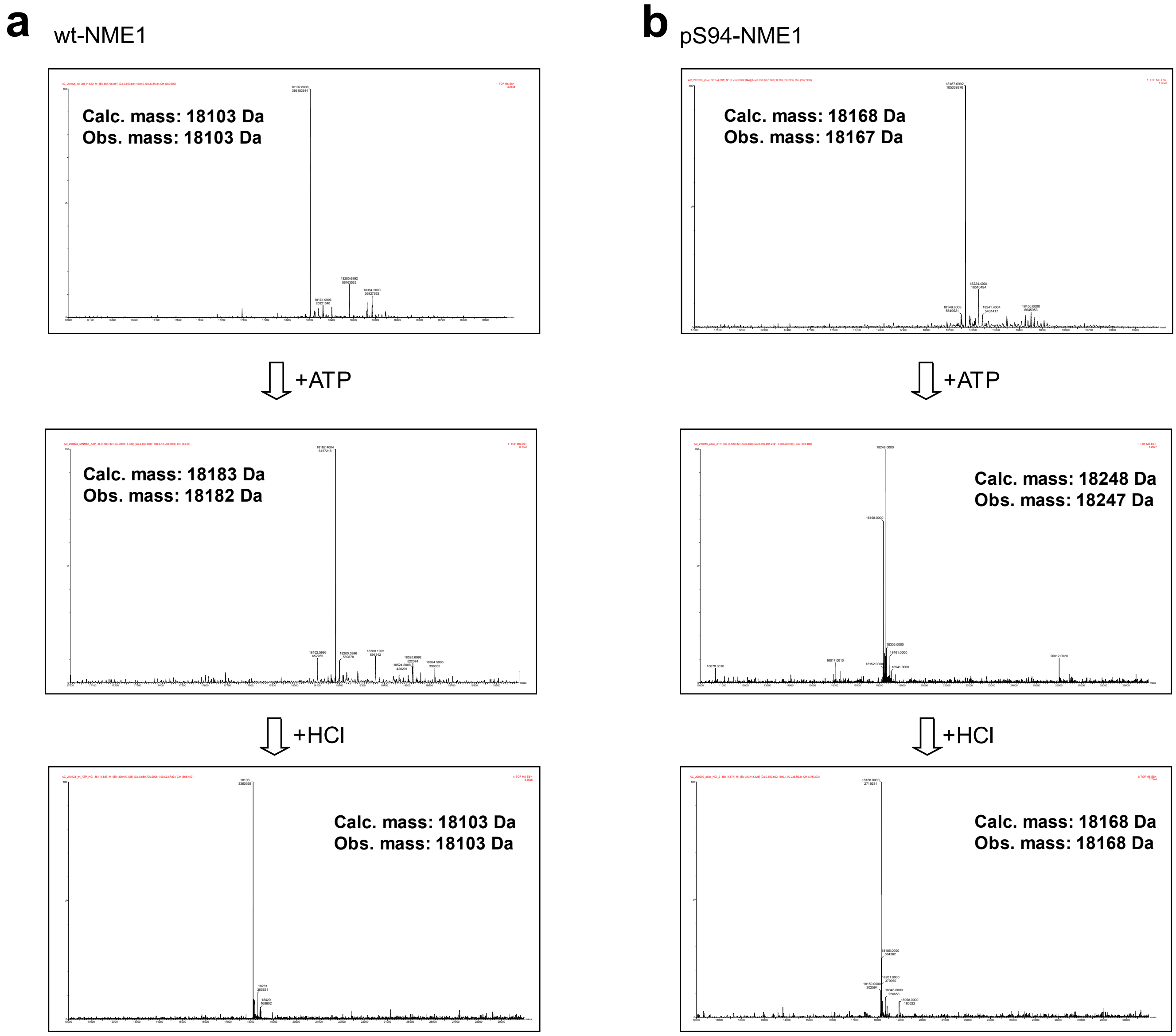


**Supplementary Figure 7.** Autophosphorylation and hydrolysis of NME1 (wt, pS94). a) wt-NME1 treated with ATP and subsequently with HCl monitored by Q-TOF-MS. b) pS94-NME1 treated with ATP and subsequently with HCl monitored by Q-TOF-MS.

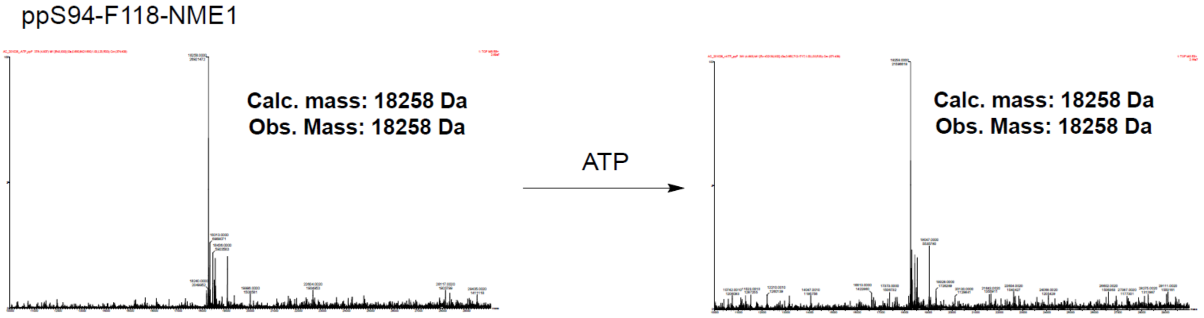


**Supplementary Figure 8.** ATP-treatment of ppS94-F118-NME1 analyzed by Q-TOF-MS. Autophosphorylation was performed at 37 °C for 1 h in 50 mM Tris-HCl, 10 mM $MgCl_2$, 1 mM DTT, and 1 mM ATP with subsequent western blot analysis using anti-NME1 and anti-1-pHis antibodies.

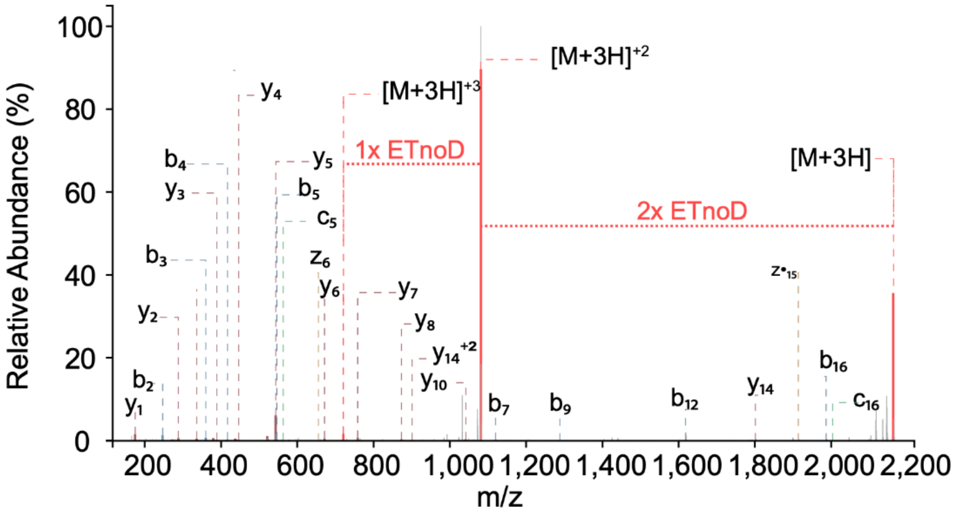


**Supplementary Figure 9.** Electron transfer no dissociation (ETnoD) event observed for the tryptic peptide containing pppS94.

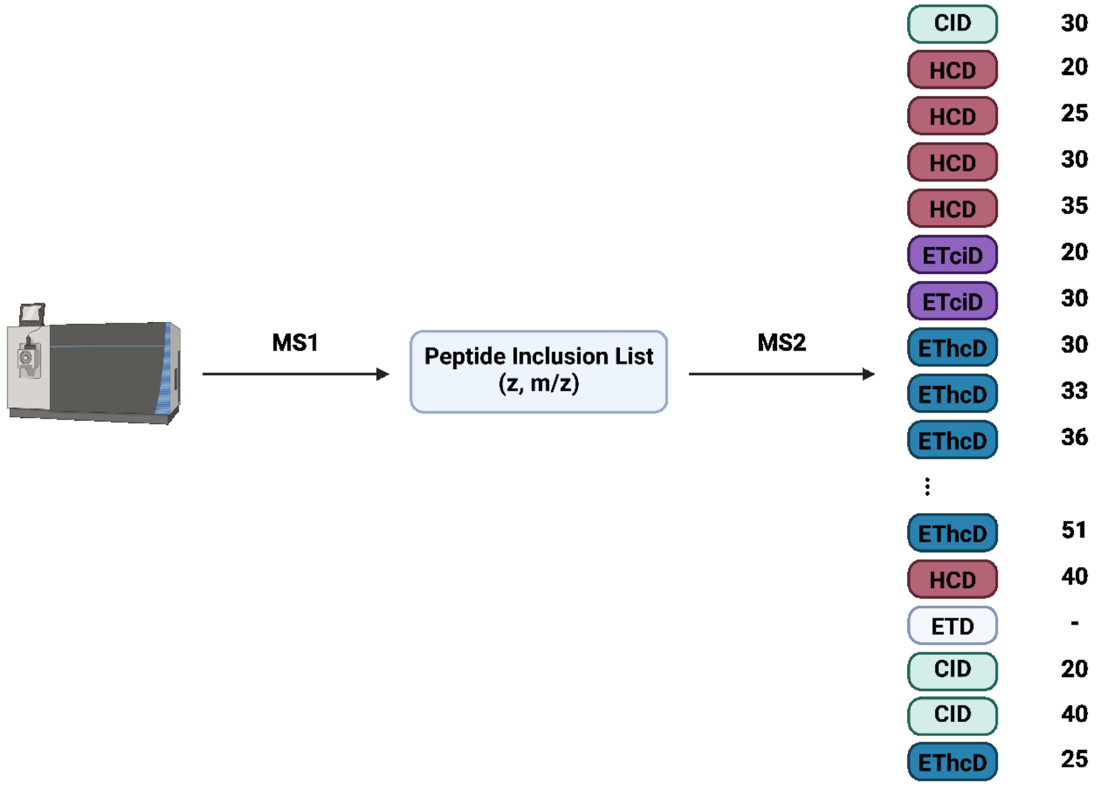


**Supplementary Figure 10.** Screening different fragmentation techniques on oligophosphorylated NME1 peptide.

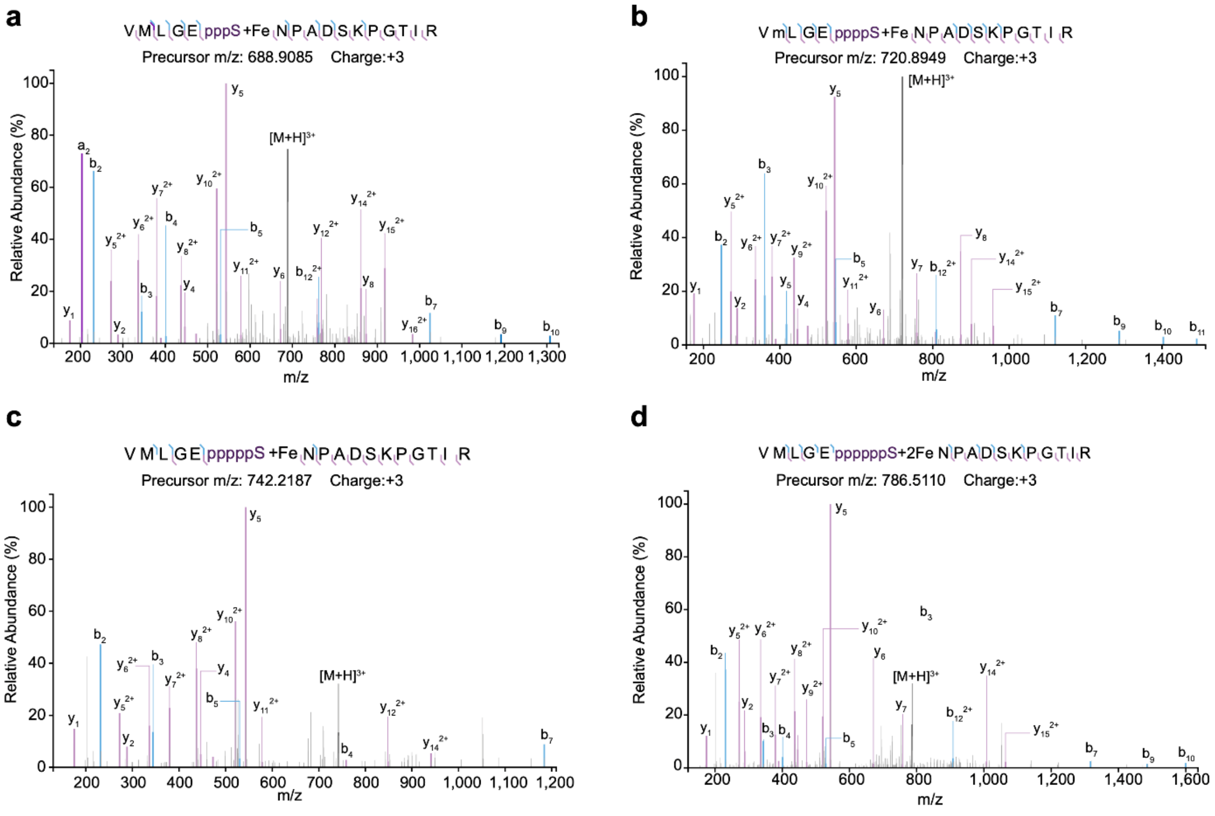

**Supplementary Figure 11.** MS/MS spectra of oligophosphorylated NME1 localizing a) triphosphorylation, b) tetraphosphorylation, c) pentaphosphorylation, and d) hexaphosphorylation on S94.

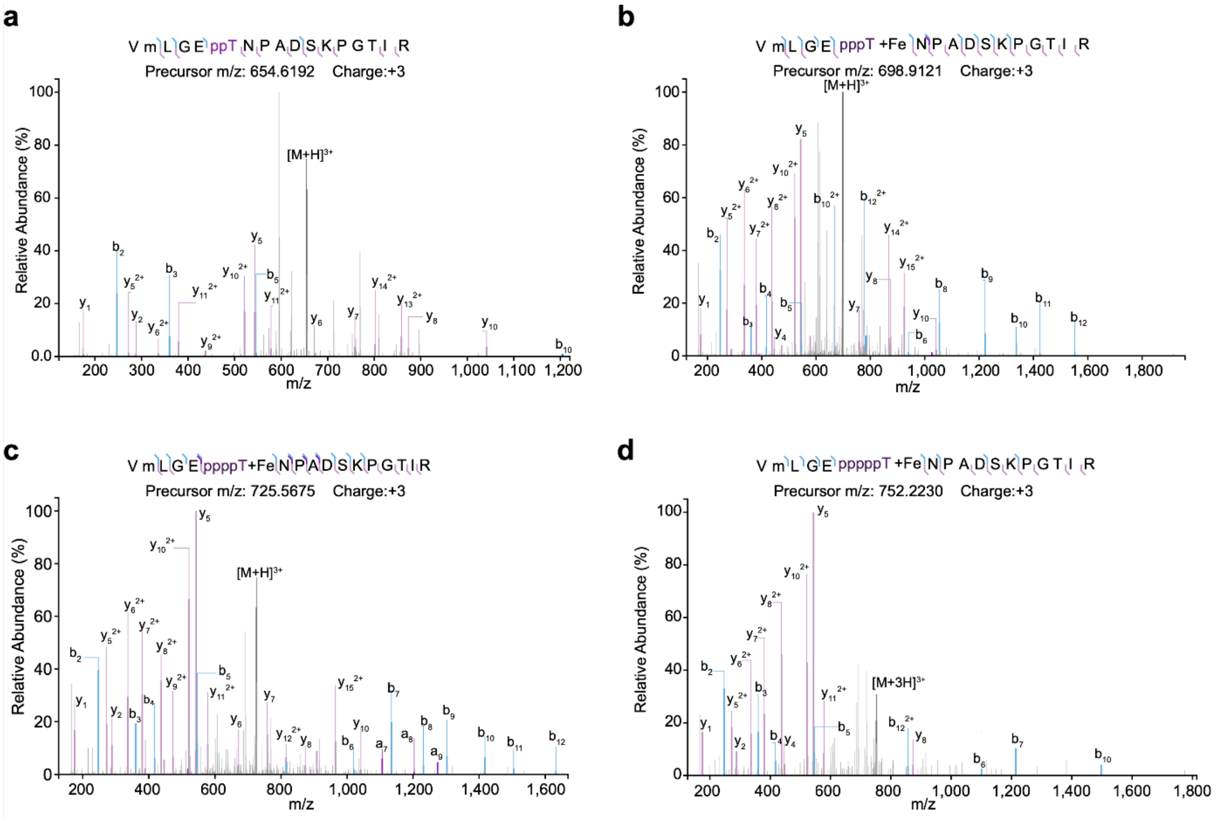


**Supplementary Figure 12.** MS/MS spectra of pyrophosphorylated and oligophosphorylated NME1 (expressed as pT94NME1) localizing a) pyrophosphorylation, b) triphosphorylation, c) tetraphosphorylation, and d) pentaphosphorylation on T94.

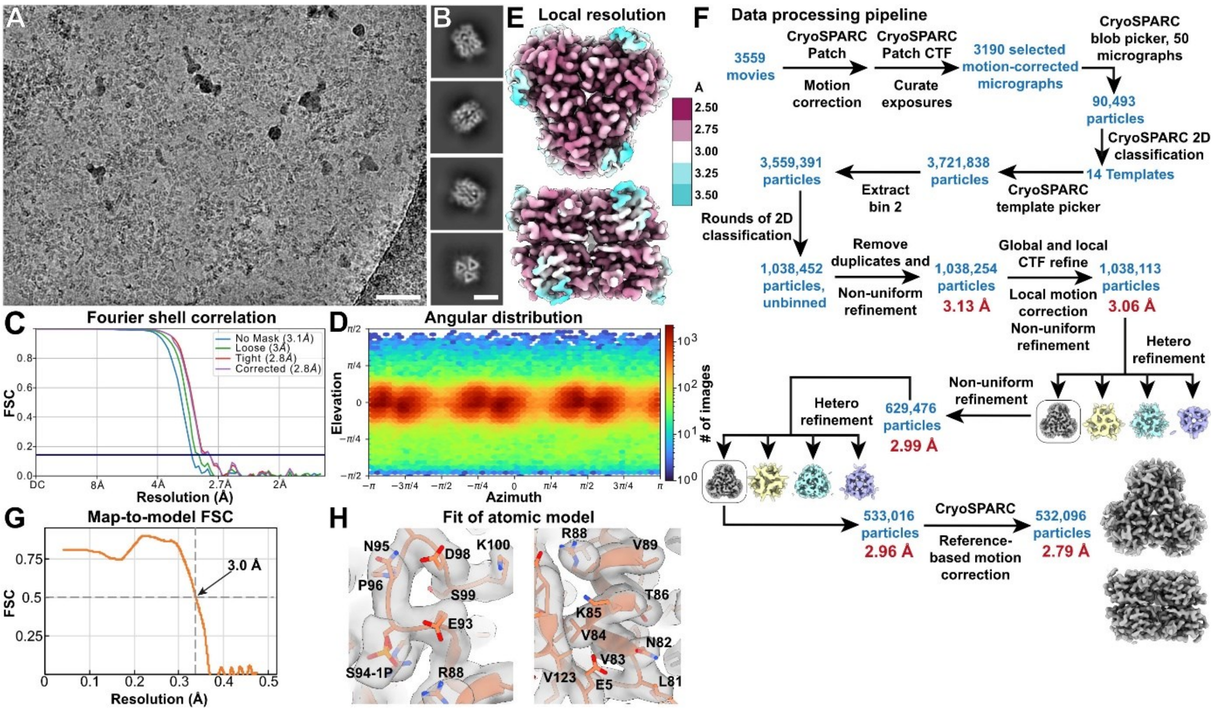


**Supplementary Figure 13: Cryo-EM and SPA of pS94-NME1.** A: Representative cryo-EM micrograph recorded at 300 kV and -1.7 µm defocus. Scale bar: 50 nm. B: Representative 2D class averages, showing hexameric pS94-NME1 in side views and top view. Scale bar: 5 nm. C,D,E: Fourier shell correlation (C), angular distribution (D), and density map colored by local resolution (E) of the final cryo-EM density map of pS94-NME1 from 532,096 particles with D3 symmetry applied. F: SPA data processing scheme. All steps were carried out in cryoSPARC, and all refinements were carried out with D3 symmetry. G: Map-to-model correlation of the pS94-NME1 model refined against the final non-sharpened density map. H: Fit of atomic model in selected parts of the density map. The density maps in E,F and H have been sharpened with DeepEMhancer.

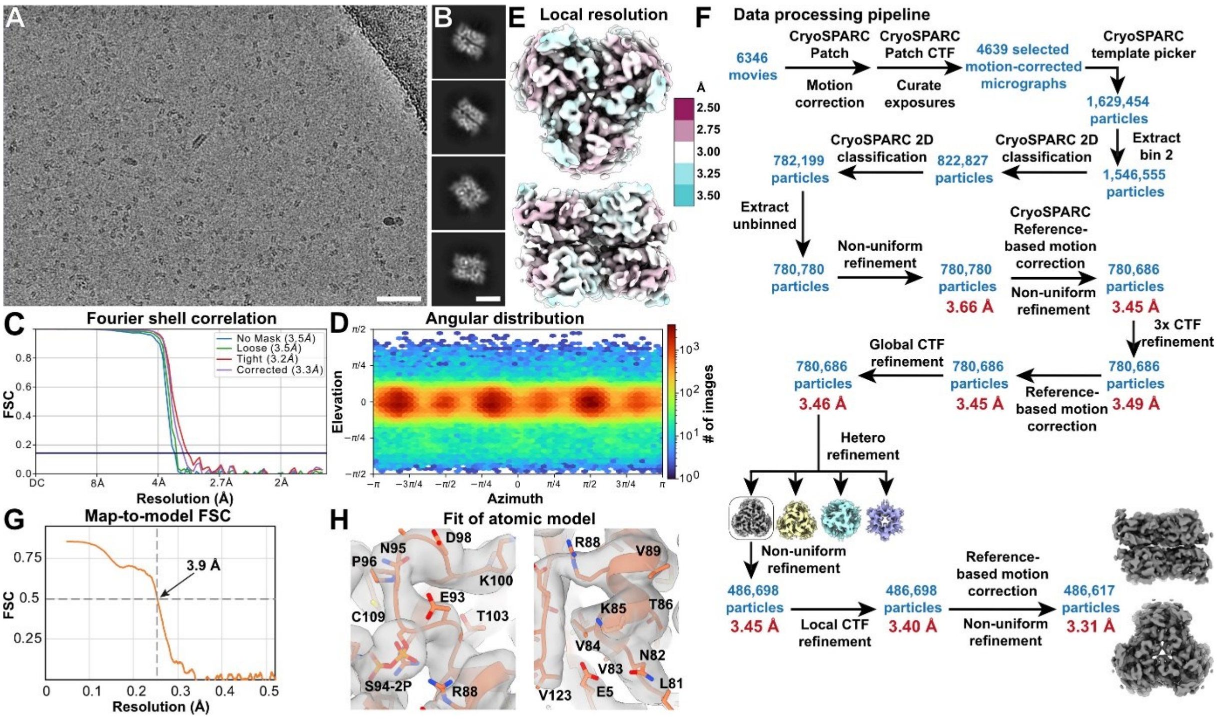


**Supplementary Figure 14: Cryo-EM and SPA of ppS94-NME1.** A: Representative cryo-EM micrograph recorded at 300 kV and -1.9 µm defocus. Scale bar: 50 nm. B: Representative 2D class averages, showing hexameric ppS94-NME1 in side views and tilted views. Scale bar: 5 nm. C,D,E: Fourier shell correlation (C), angular distribution (D), and density map colored by local resolution (E) of the final cryo-EM density map of ppS94-NME1 from 486,698 particles with D3 symmetry applied. F: SPA data processing scheme. All steps were carried out in cryoSPARC, and all refinements were carried out with D3 symmetry. G: Map-to-model correlation of the ppS94-NME1 model refined against the final non-sharpened density map. H: Fit of atomic model in selected parts of the density map. The density maps in E,F and H have been sharpened with DeepEMhancer.

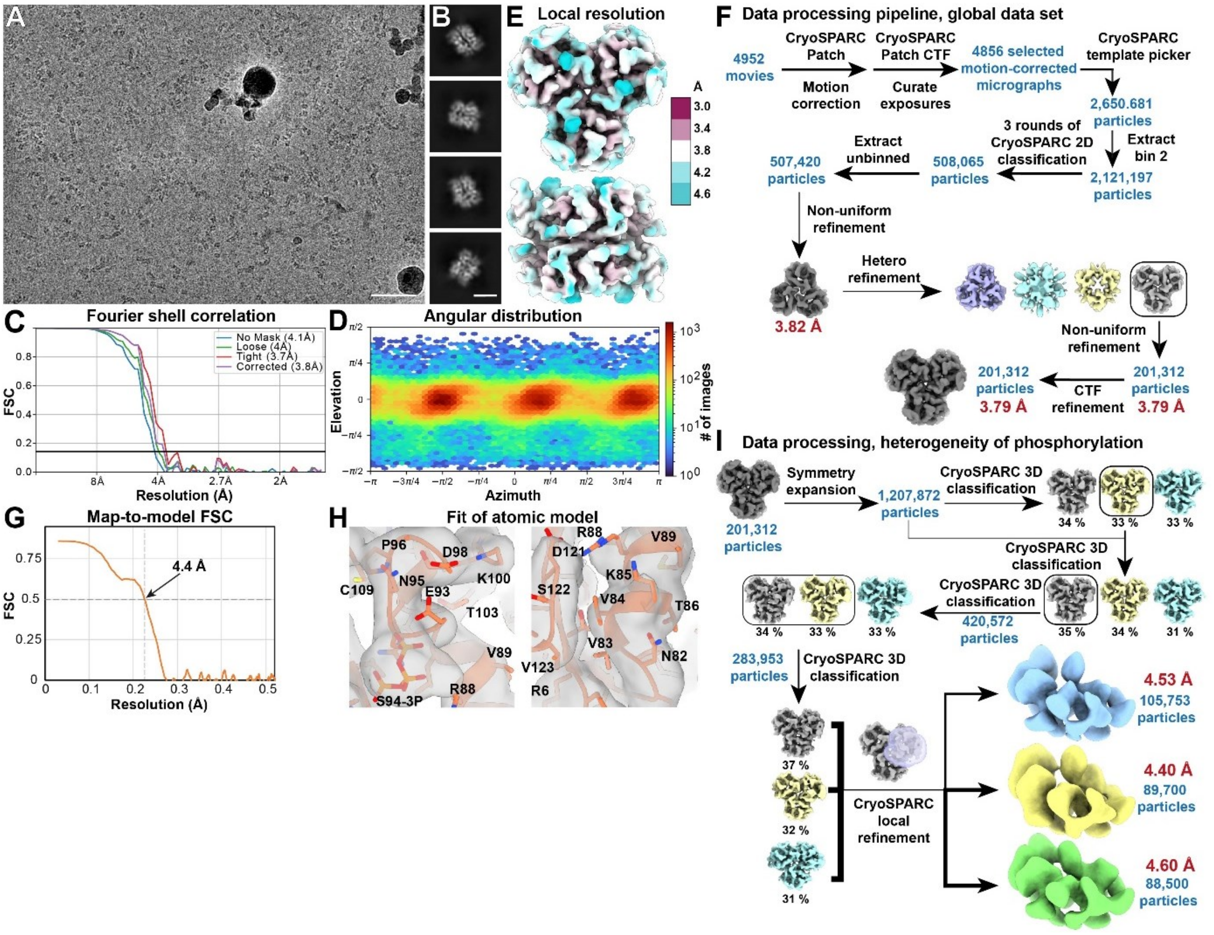


**Supplementary Figure 15: Cryo-EM and SPA of Oligo-pS94-NME1.** A: Representative cryo-EM micrograph of Oligo-pS94-NME1 (i.e., ppS94-NME1 with 1mM ATP) recorded at 300 kV and -1.9 µm defocus. Scale bar: 50 nm. B: Representative 2D class averages, showing hexameric Oligo-pS94-NME1 in side views and tilted views. Scale bar: 5 nm. C,D,E: Fourier shell correlation (C), angular distribution (D), and density map colored by local resolution (E) of the final cryo-EM density map of Oligo-pS94-NME1 from xxx particles with D3 symmetry applied. F: SPA data processing scheme applied for the global map with D3 symmetry. All steps were carried out in cryoSPARC, and all refinements were carried out with D3 symmetry. G: Map-to-model correlation of the pppS94-NME1 model refined against the final non-sharpened density map. H: Fit of atomic model in selected parts of the density map. I: Data processing scheme after symmetry expansion to identify heterogeneity in phosphorylation. The density maps in E,F and H have been sharpened with DeepEMhancer.

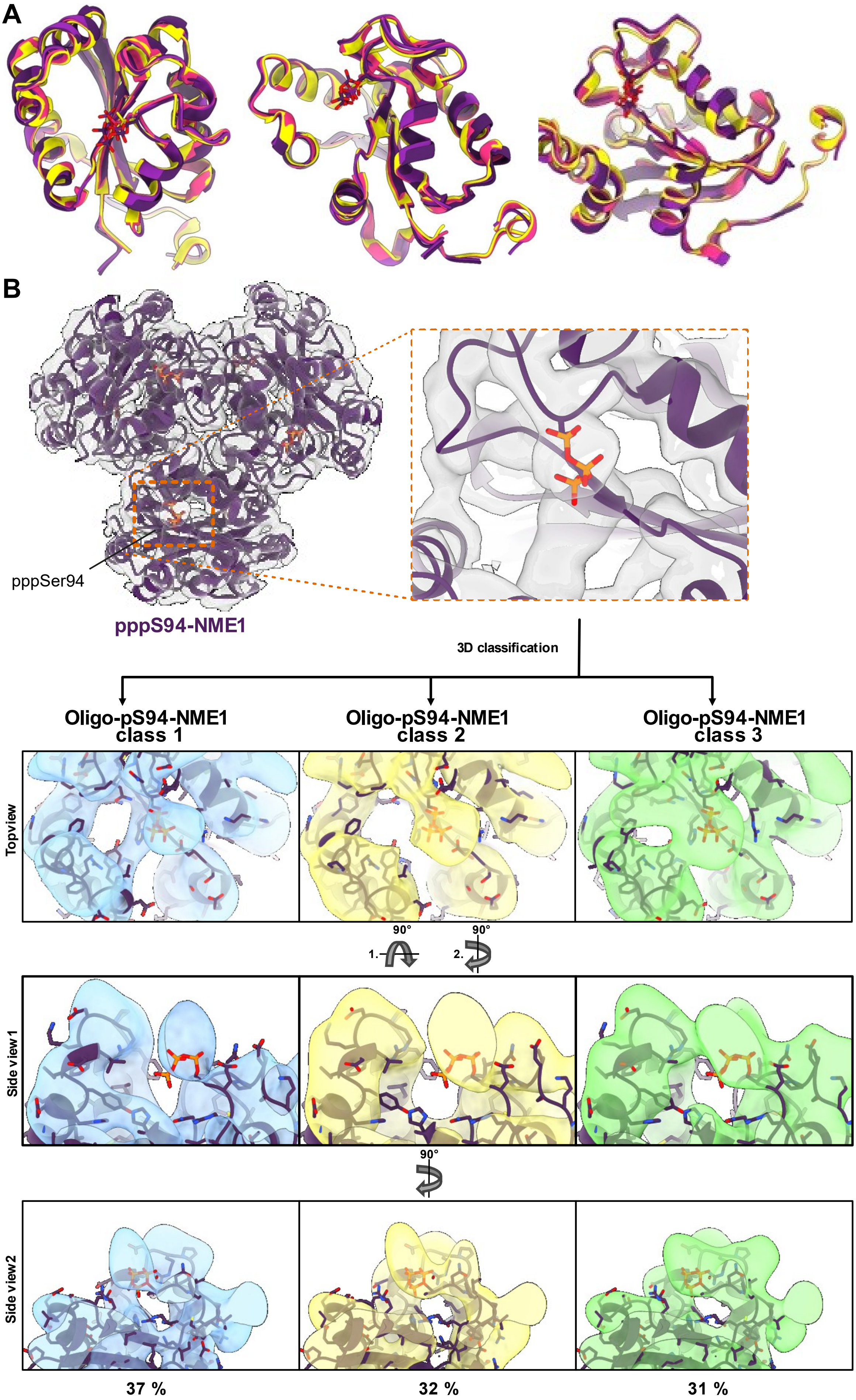
A
B
pppSer94
pppS94-NME1
3D classification
Oligo-pS94-NME1 class 1
Oligo-pS94-NME1 class 2
Oligo-pS94-NME1 class 3
Topview
90°
1.
2.
Side view1
Side view2
37 %
32 %
31 %

**Supplementary Figure 16**. Symmetry expansion and focused 3D classification of one subunit. a) Overlay of one subunit of wt-NME1, pS94-NME1, ppS94-NME1, and oligopS94-NME1. b) Identification of three subsets with different extends of additional cryo-EM density appearing at the protein’s surface shown by a top view and side view.

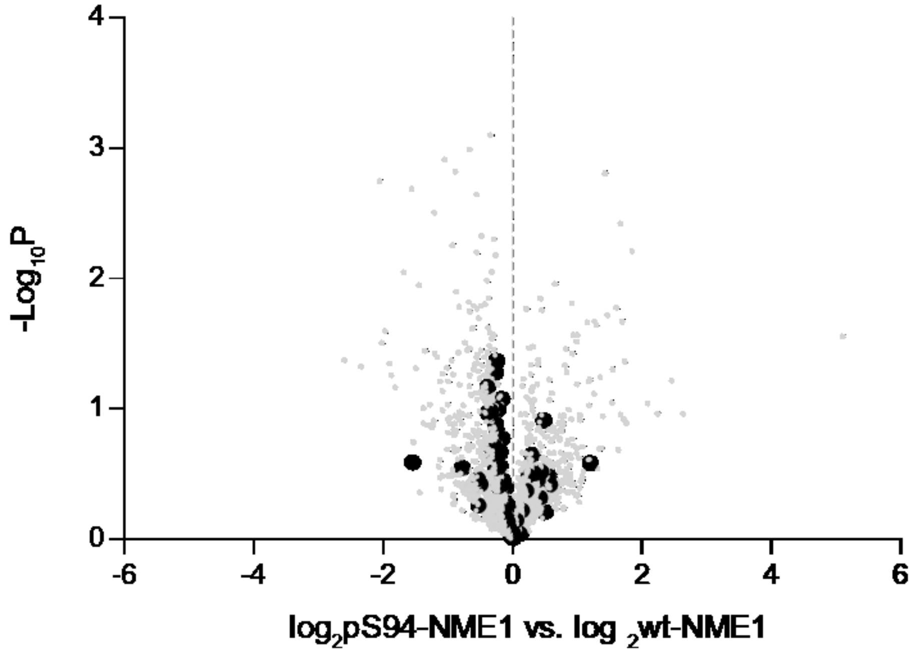


**Supplementary Figure 17**. Volcano plot depicting LFQ values of pS94-NME1 versus wt-NME1 after a t-test. The x-axis display the difference of LFQ values on a $\log_2$ scale and the y-axis shows the $-\log_{10}P$ value. Labeled hits (large black dots) represents known interactors (BioGRID database) prefentially enriched with wt-NME1 or pS94-NME1.

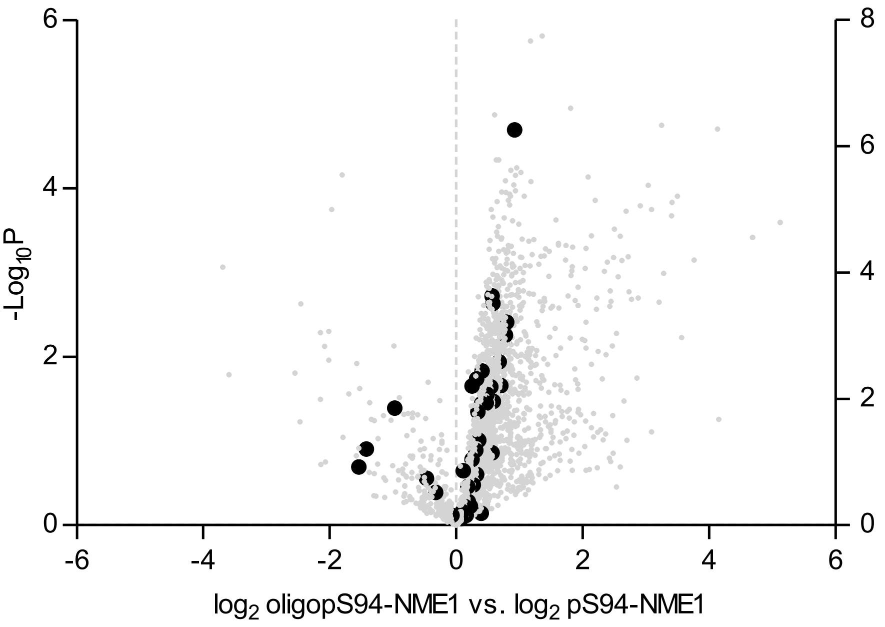


**Supplementary Figure 18**. Volcano plot depicting LFQ values of oligopS94-NME1 versus pS94-NME1 after a t-test. The x-axis display the difference of LFQ values on a $log_2$ scale and the y-axis shows the $-log_{10}P$ value. Labeled hits (large black dots) represents known interactors (BioGRID database) prefentially enriched with pS94-NME1 or oligopS94-NME1.

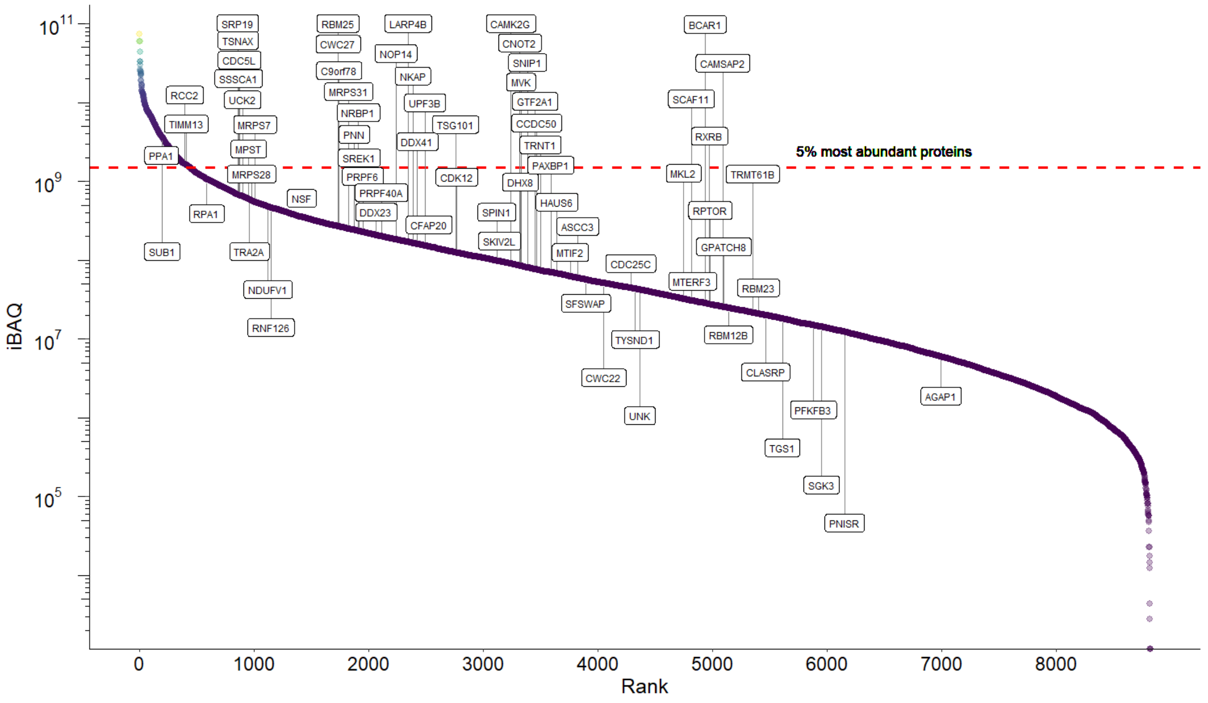


**Supplementary Figure 19**. iBAQ values of a whole HEK293T proteome were ranked and plotted against the $log_{10}$ iBAQ values. Protein interactors of oligopS94-NME1 (vs wt-NME1 and pS94-NME1) were highlighted.

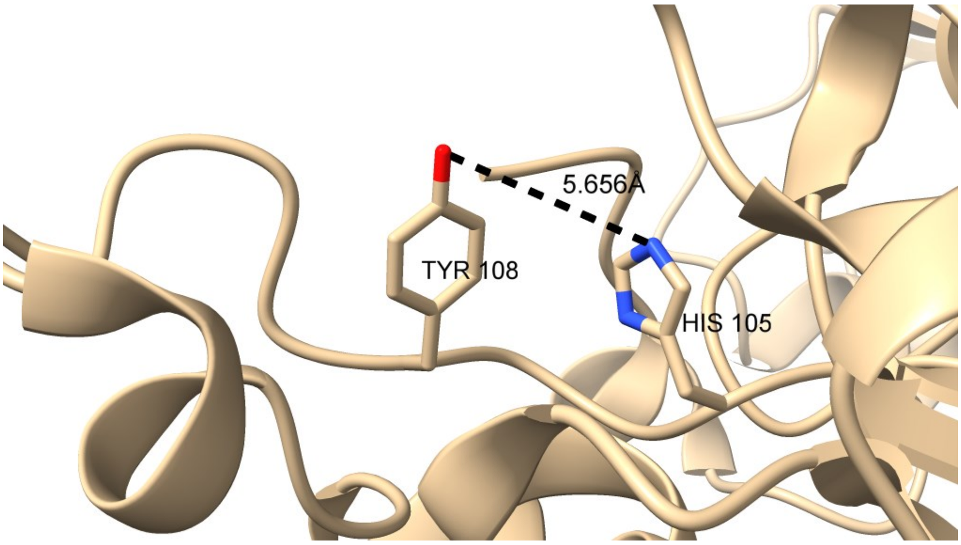


**Supplementary Figure 20.** Crystal structure of PGAM5 highlighting the location of H105 and Tyr108 (PDB code: 3MXO).

#### Extended Methods

##### Q-TOF-MS

High-resolution ESI-MS spectra were recorded on two different instruments: 1) Agilent 6220 TOF Accurate Mass coupled to an Agilent 1200 LC (Agilent Technologies, USA) and were measured at 35 ˚C between 100–2000 m/z. The used column was an Accucore RP-MS (30 x 2.1 mm; 2.6 µm particle size) eluted with a flow of 0.8 mL/min and the following gradient (A = $H_2O$ + 0.1% TFA, B = MeCN + 0.1% TFA), gradient: 5% B 0–0.2 min, 5–99% B 0.2–1.1 min, 99% B 1.1–2.5 min. 2) Agilent Technologies 6230 Accurate Mass TOF LC/MS linked to Agilent Technologies HPLC 1260 Series; Column: Thermo Accucore RP-MS; Particle Size: 2.6 µM Dimension: 30 x 2.1 mm. The following gradient was used: A = $H_2O$ + 0.1 % formic acid, B = MeCN + 0.1 % formic acid, 5% B 0.0–0.2 min, 5-99% B 0.2–1.1 min, 99% B 1.1–3.6 min, 5% B 3.6–4.9 min. Flow rate: 0.8 mL/min; UV-detection: 220 nm, 254 nm, 300 nm.

##### Intact protein MS

Intact proteins were analyzed using a Waters H-class instrument equipped with a quaternary solvent manager, a Waters sample manager-FTN, a Waters PDA detector, and a Waters column manager with an Acquity UPLC protein BEH C4 column (300 Å, 1.7 µm, 2.1 mm x 50 mm). Proteins were eluted at a column temperature of 80 ˚C with a flow rate of 0.3 mL/min. The following gradient was used: A = $H_2O$ + 0.01% formic acid, B = MeCN + 0.01% formic acid. 5–95% B 0–6 min at 40 ˚C. Mass analysis was conducted with a Waters XEVO G2-XS Q-TOF analyzer. Proteins were ionized in positive ion mode applying a cone voltage of 40 kV.

***Cryo-EM sample preparation and data acquisition.***

For pS94-NME1 and ppS94-NME1, purified protein was vitrified on Quantifoil 1.2/1.3 Cu 300 mesh grids at 0.9 mg/ml using a Vitrobot Mark IV set to a blot force of 0, blotting time of 3.0 For oligo-pS94-NME1, ppS94-NME1 was incubated at 1 mM concentration with ATP for 3 h at 37˚C before vitrification as above. Micrographs for all three data sets were acquired using a FEI Titan Krios G3i microscope (Thermo Fisher Scientific) operated at 300 kV equipped with a Bioquantum K3 direct electron detector and energy filter (Gatan) running in CDS counting mode at a slit width of 20 eV and at a nominal magnification of 105,000×, giving a calibrated physical pixel size of 0.83 Å/px on the specimen level. EPU 2.12 was utilized for automated data acquisition with AFIS enabled. For pS94-NME1, movies were recorded for 2.0s accumulating a total electron dose of 47.7 $e^{-}/Å^{2}$ fractionated into 50 frames. Nominal defocus values were between -1.1 and -2.6 µm. For ppS94-NME1, movies were recorded for 2.0 s resulting in a total electron dose of 44.6 $e^{-}/Å^{2}$ fractionated into 50 frames with nominal defocus values between -1.2 and -2.4. For NME1-94Ser-Oligo-P, movies were recorded for 2.0 s accumulating a total electron dose of 44.1 $e^{-}/Å^{2}$ distributed over 50 frames. Here, nominal defocus values were between -1.4 and -2.4.

***Data processing of pS94-NME1***

All data processing steps were carried out using CryoSPARC and are outlined in **Supplementary Fig. S12F**.[1] 3555 movies were aligned using Patch Motion correction and CTF was determined using patch CTF estimation. After sorting out bad images, 3190 micrographs were selected for further processing. 90,493 particles were selected from 50 micrographs using the blob picker and extracted for initial 2D classification and generation of autopicking templates. Subsequent template-based autopicking using 14 selected 2D classes as templates and particle curation identified 3,721,838 particles. Of these, 3,559,391 were extracted (2x binning) for 2D classification. After three rounds of 2D classification (70 online-EM interations), an initial 3D hetero refinement (3 classes), and duplicate removal, 1,038,254 particles were re-extracted without binning and subjected to non-uniform refinement using D3 symmetry and a molecular map of PDB 5UI4 (lowpass filtered to 12 Å) as an initial model.[2] After iterative rounds of global and local CTF refinement and local motion correction, 1,038,113 particles were sorted in two successive rounds of 3D hetero refinement (4 classes each), resulting in a final class of 533,016 particles.[3] These were then further processed using non-uniform refinement, followed by reference-based motion correction, yielding 532,096 particles. A final non-uniform refinement resulted in a resolution of 2.8 Å according to the gold-standard Fourier Shell Correlation (FSC) criterion **Supplementary Fig. S12C, E**. DeepEMhancer[4] was applied for map sharpening.

***Data processing of ppS94-NME1.***

All data processing steps were performed using CryoSPARC and are outlined in Figure **Supplementary Fig. S13F**.[1] 6346 movies were aligned using patch motion correction and patch CTF estimation was used to determine the CTF. After rejecting bad images, 4639 micrographs were selected for further processing and subjected to particle picking using the template picker, which identified 1,629,454 particles. Particles were extracted with 2x binning, resulting in 1,546,555 particles that were sorted using two successive 2D classifications (70 online-EM iterations). 2D classes were selected and 782,199 corresponding particles were extracted (no binning), resulting in 780,780 particles that were subjected to non-uniform refinement using 3D symmetry and the pS94-NME1 map as the initial model.[2] After iterative rounds of global and local CTF refinement and local motion correction, 780,686 particles were sorted using heterogenous refinement.[3] One class with 486,698 particles was selected and further refined using non-uniform refinement, which was followed by a local CTF refinement and reference-based motion correction. The resulting 486,617 particles were refined with non-uniform refinement, resulting in a resolution of 3.31 Å according to the gold-standard Fourier Shell Correlation (FSC) criterion **Supplementary Fig. S13C, E**. DeepEMhancer[4] was used for map sharpening.

***Data processing of oligo-pS94-NME1***

All data processing steps were carried out using CryoSPARC and are outlined in **Supplementary Fig. S14F, I**.[1] 4952 movies were aligned using Patch Motion correction and the CTF was determined using patch CTF estimation. After rejecting bad images, 4856 micrographs were selected for further processing. Particles were picked using the template picker, identifying 2,650,681 particles, which were extracted using 2x binning, retaining 2,121,197 particles. Sorting was performed using three consecutive rounds of 2D classification (70 online-EM iterations), resulting in a total of 508,065 total particles. After extraction (no binning) 507,420 particles were retained, subjected to non-uniform refinement with D3 symmetry using the ppS94-NME1 map as initial model and further sorted using heterogeneous refinement.[2] The class containing 201,312 particles was selected and refined using non-uniform refinement, resulting in a resolution of 3.79 Å according to the gold-standard Fourier Shell Correlation (FSC) criterion **Supplementary Fig. S14C, E**. DeepEMhancer[4] was used for map sharpening. To investigate the heterogeneity of S94 phosphorylation the particles were symmetry expanded, resulting in 1,207,872 particles that were subjected to four consecutive 3D classifications using a mask focused on one NME1 monomer and forced hard classification. Three classes representing 283,953 particles were each subjected to local refinement using a focus mask on one NME1 monomer, resulting three maps of 4.53 Å (105,753 particles), 4.40 Å (89,700 particles) and 4.60 Å (88,500 particles) according to the gold-standard Fourier Shell Correlation (FSC) criterion (**Supplementary Fig. S14I**).

***Atomic modeling of pS94-NME1***

The atomic model of human NME1 (PDB 5UI4) was used as starting model and the attached imidazole fluorosulfate group and water molecules were removed. The model was rigid-body fitted into the sharpened density map using UCSF ChimeraX[5], manually adjusted in Coot[6], where Thr94 was mutated to phosphoserine, and ISOLDE[7], and then refined using real-space refinement in Phenix[8] (**Supplementary Fig. S12G, H**). Cryo-EM data processing and model refinement statistics are summarized in Table S1.

***Atomic modeling of ppS94-NME1 and pppS94-NME1***

The atomic model of pS94-NME1 (see above) was used as starting point. The model was first rigid-body fitted into the obtained sharpened density maps for ppS94-NME1and oligo-pS94-NME1, respectively, using UCSF ChimeraX[5] and then manually adjusted in Coot. Geometry retraints for ppS and pppS were generated using phenix.elbow[9], and the resulting cif files were manually changed from a ligand to an amino acid. The changed amino acids were incorporated in the models, manually adjusted in Coot, and then refined using real-space refinement in Phenix[8] (**Supplementary Fig. S13 and S14G, H**). Cryo-EM data processing and model refinement statistics are summarized in Table S1.

#### Chemical Synthesis and Characterization

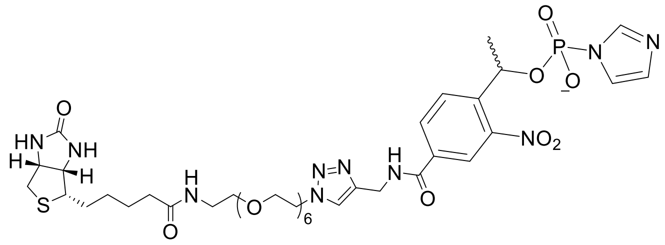

Biotin-PEG$_6$-triazole-NPE-(1H-imidazolide-1-yl)phosphonate was synthesized as previously described.[10] Spectral data matches previously reported values.

### Q-TOF-Spectra

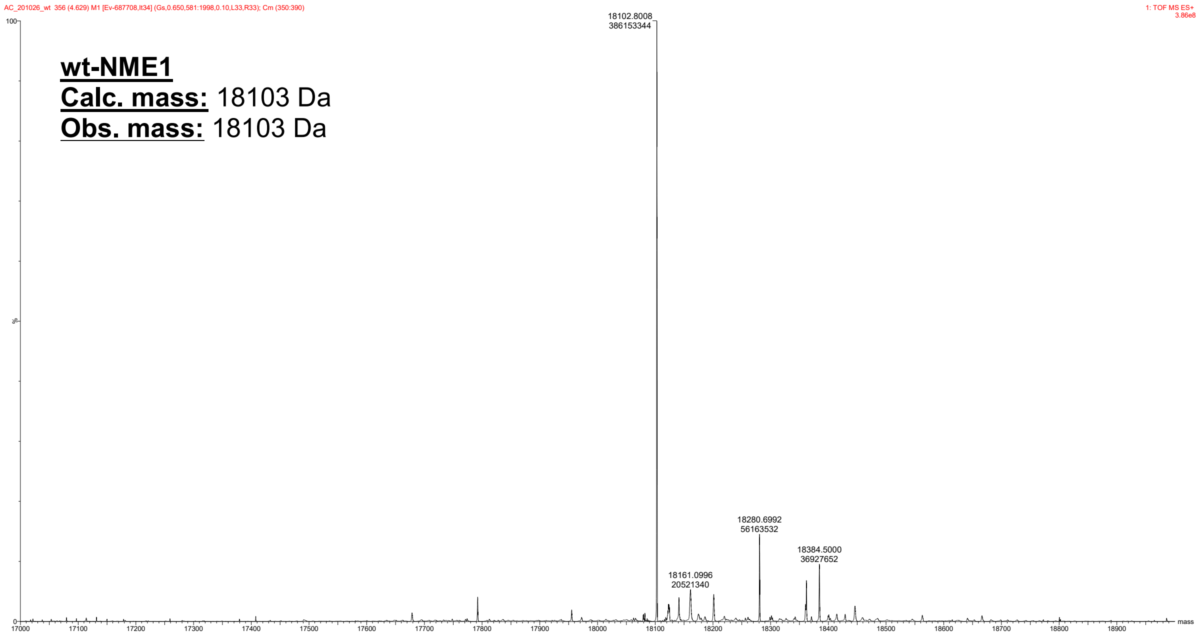


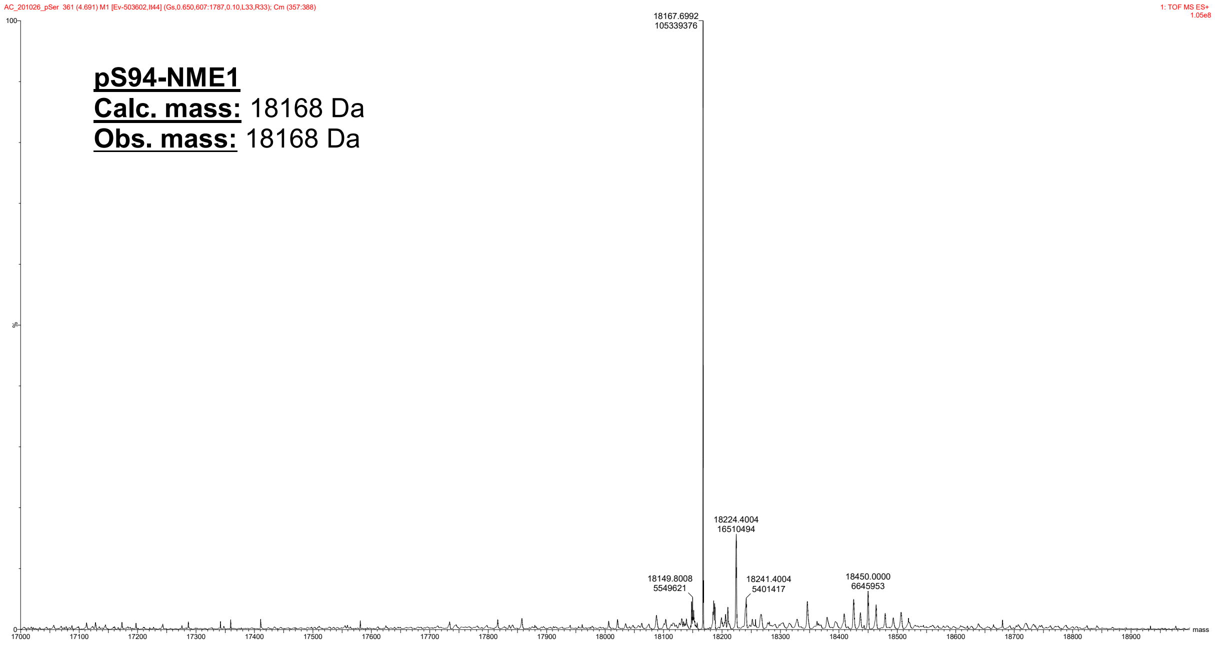

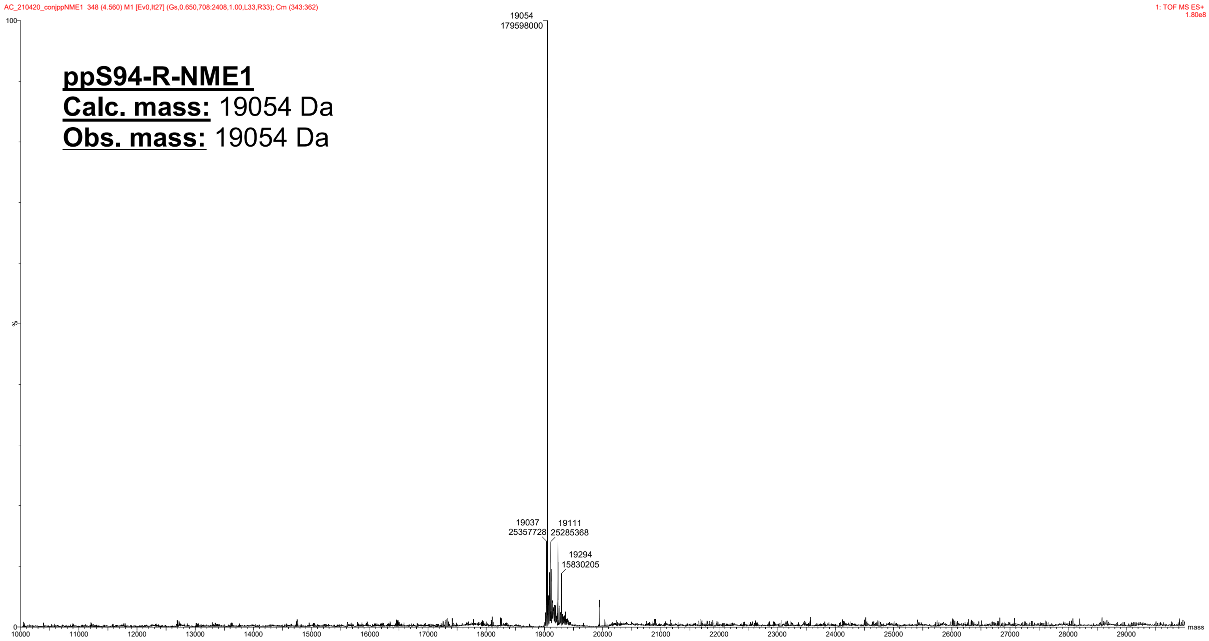
ppS94-R-NME1
Calc. mass: 19054 Da
Obs. mass: 19054 Da
19054
179598000
19037
25357728
19111
25285368
19294
15830205
1: TOF MS ES+
1.80e8
mass

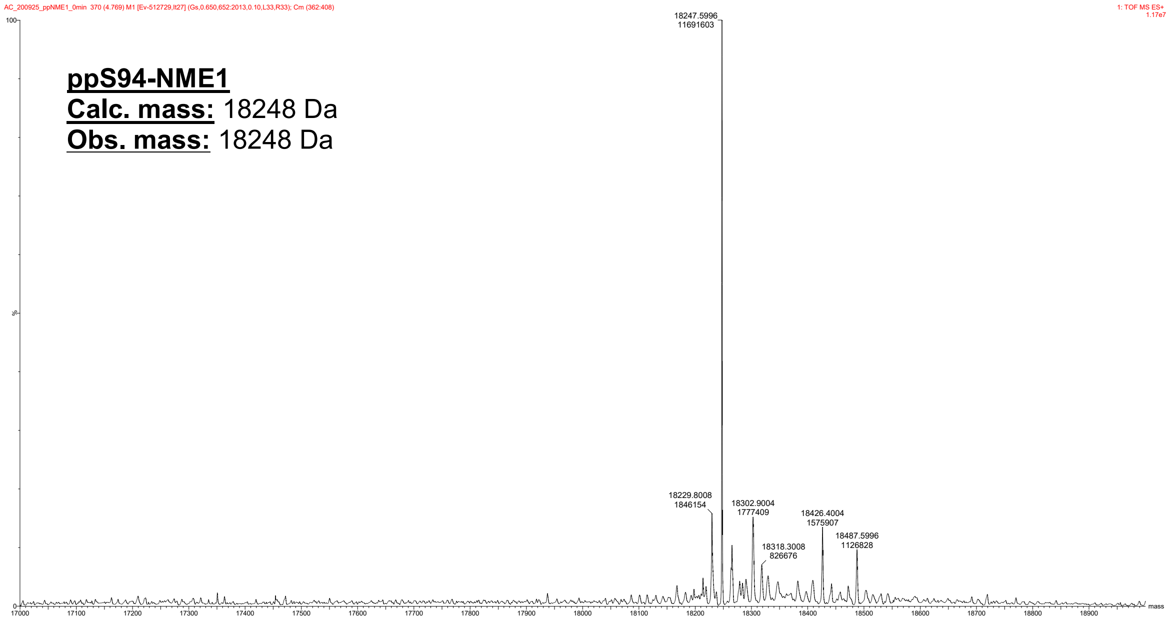
ppS94-NME1
Calc. mass: 18248 Da
Obs. mass: 18248 Da
18247.5996
11691603
18229.8008
1846154
18302.9004
1777409
18426.4004
1575907
18487.5996
1126828
18318.3008
826676
1: TOF MS ES+
1.17e7
mass

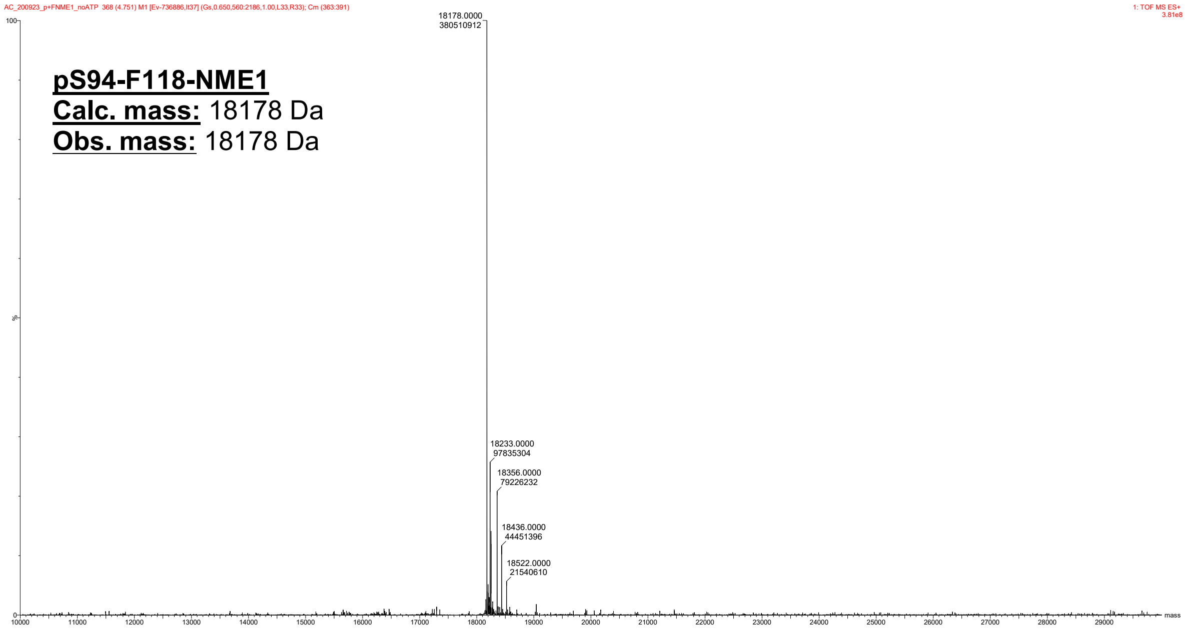
AC_200923_p+FNME1_noATP 368 (4.751) M1 [Ev-736886,It37] (Gs,0.650,560:2186,1.00,L33,R33); Cm (363:391)
1: TOF MS ES+
3.81e8
pS94-F118-NME1
Calc. mass: 18178 Da
Obs. mass: 18178 Da
18178.0000
380510912
18233.0000
97835304
18356.0000
79226232
18436.0000
44451396
18522.0000
21540610
mass

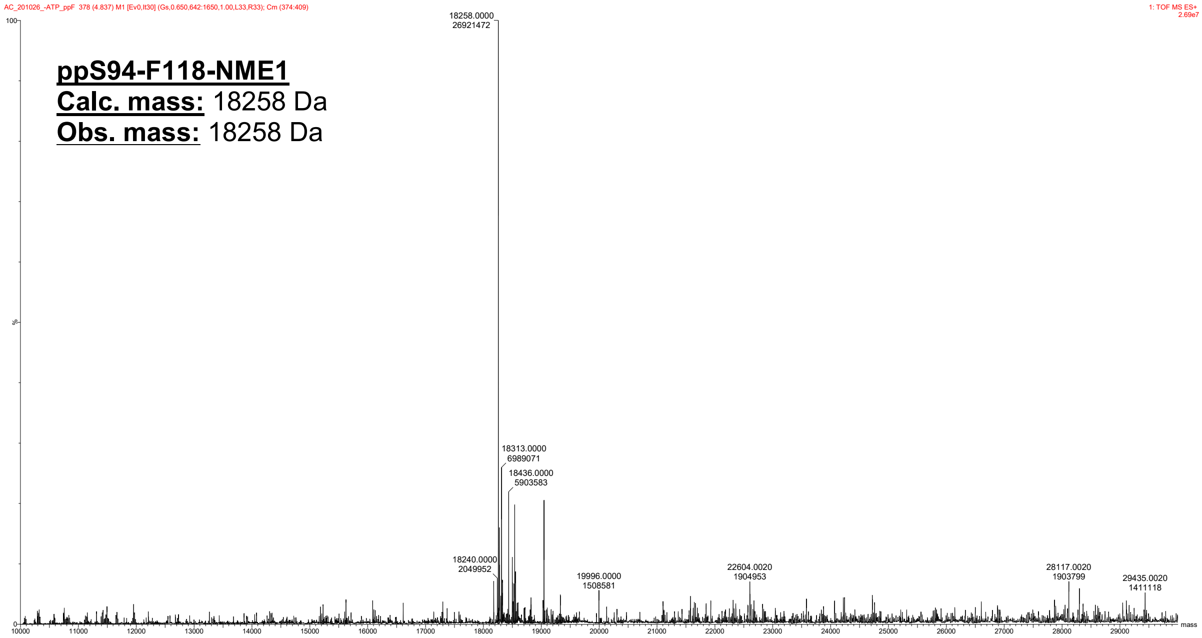
AC_201026_-ATP_ppF 378 (4.837) M1 [Ev0,It30] (Gs,0.650,642:1650,1.00,L33,R33); Cm (374:409)
1: TOF MS ES+
2.69e7
ppS94-F118-NME1
Calc. mass: 18258 Da
Obs. mass: 18258 Da
18258.0000
26921472
18313.0000
6989071
18436.0000
5903583
18240.0000
2049952
19996.0000
1508581
22604.0020
1904953
28117.0020
1903799
29435.0020
1411118
mass

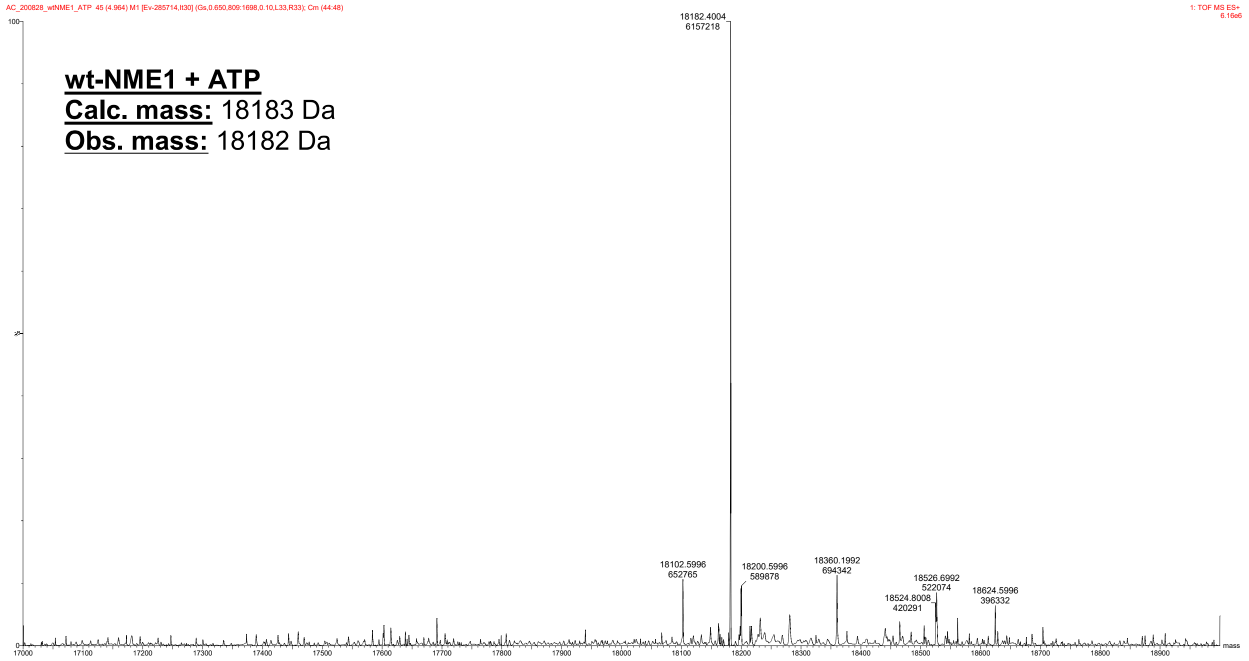
AC_200828_wtNME1_ATP 45 (4.964) M1 [Ev-285714,It30] (Gs,0.650,809:1698,0.10,L33,R33); Cm (44:48)
1: TOF MS ES+
6.16e6
wt-NME1 + ATP
Calc. mass: 18183 Da
Obs. mass: 18182 Da
18182.4004
6157218
18102.5996
652765
18200.5996
589878
18360.1992
694342
18526.6992
522074
18524.8008
420291
18624.5996
396332
mass

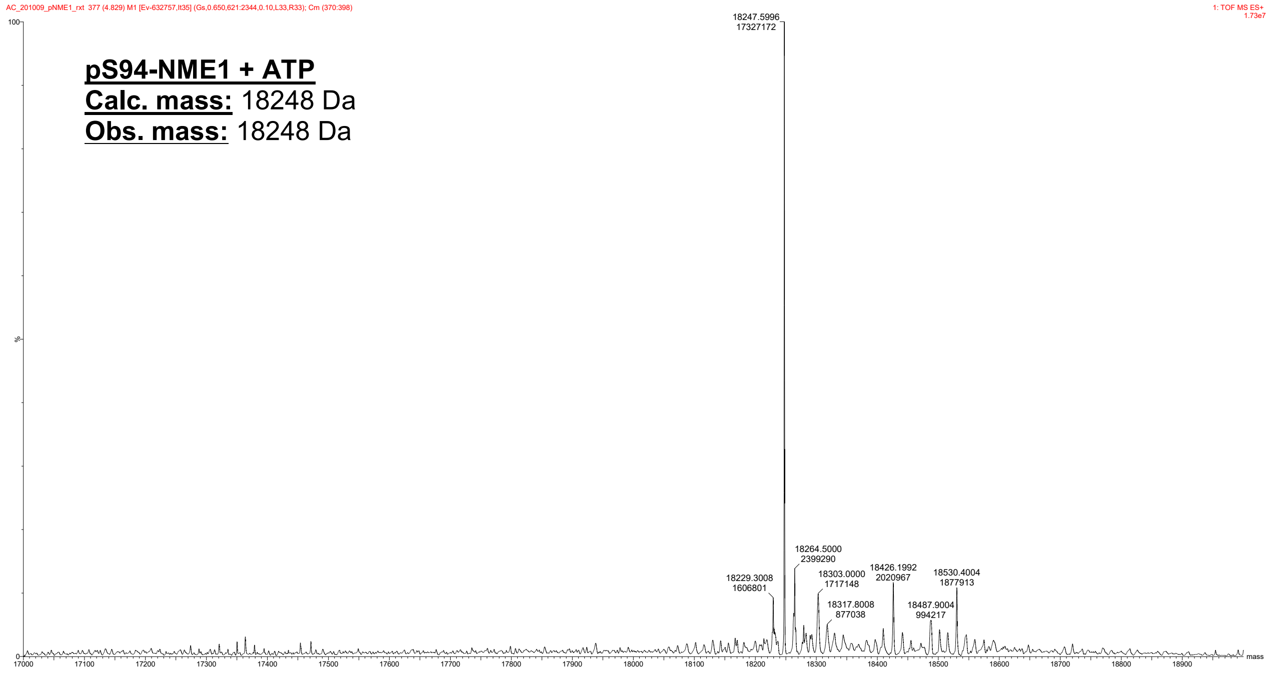
AC_201009_pNME1_rxt 377 (4.829) M1 [Ev-632757,It35] (Gs,0.650,621:2344,0.10,L33,R33); Cm (370:398)
1: TOF MS ES+
1.73e7
pS94-NME1 + ATP
Calc. mass: 18248 Da
Obs. mass: 18248 Da
18247.5996
17327172
18264.5000
2399290
18229.3008
1606801
18303.0000
1717148
18426.1992
2020967
18530.4004
1877913
18317.8008
877038
18487.9004
994217
mass

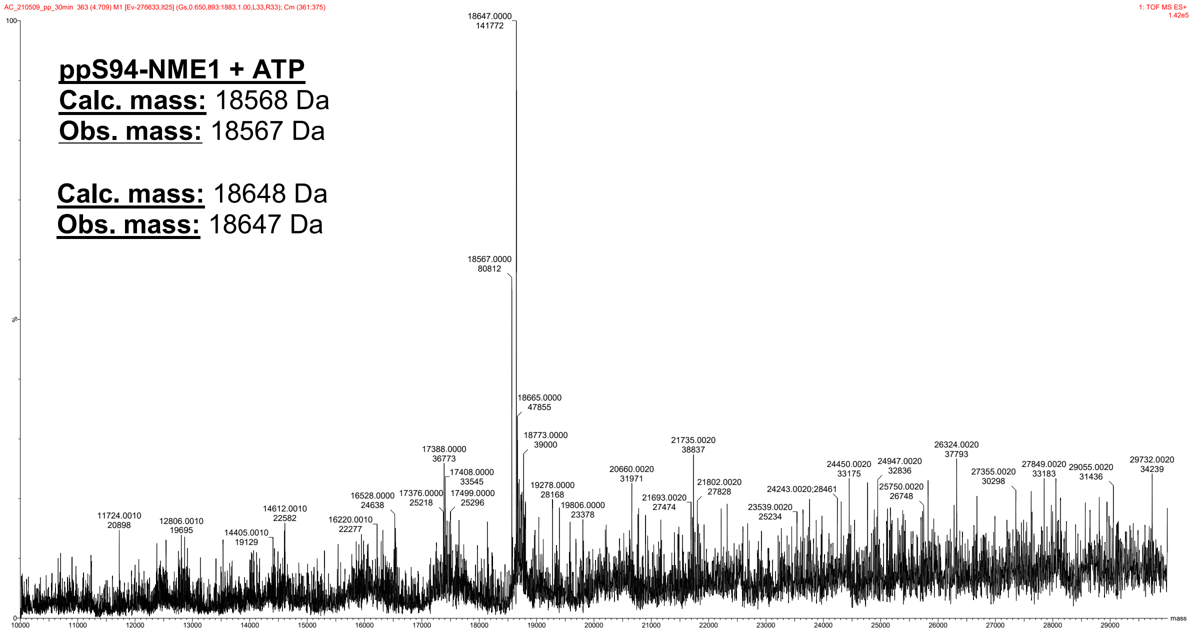
ppS94-NME1 + ATP
Calc. mass: 18568 Da
Obs. mass: 18567 Da
Calc. mass: 18648 Da
Obs. mass: 18647 Da

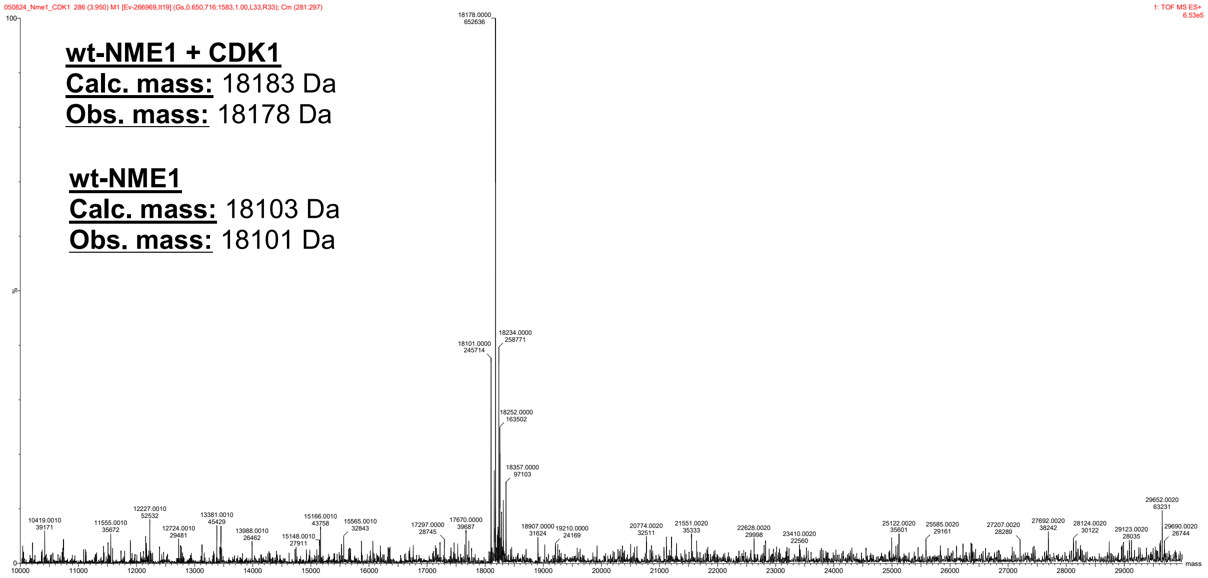
wt-NME1 + CDK1
Calc. mass: 18183 Da
Obs. mass: 18178 Da
wt-NME1
Calc. mass: 18103 Da
Obs. mass: 18101 Da

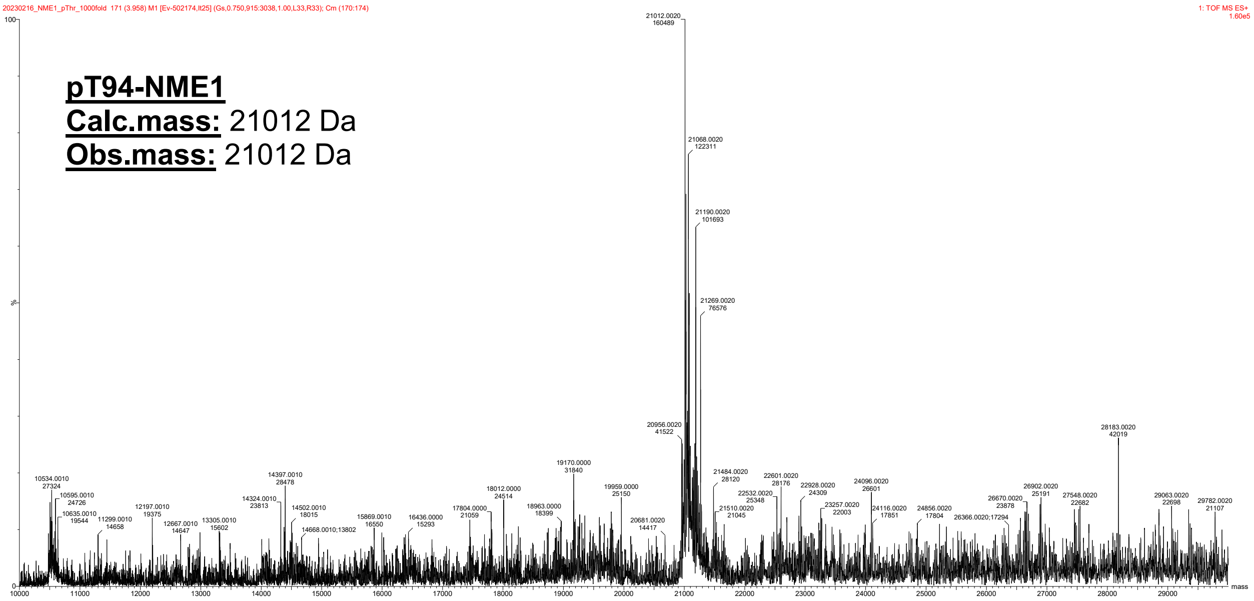
